# A subset of dorsal raphe dopamine neurons is critical for survival-oriented vigilance

**DOI:** 10.1101/2025.03.31.646345

**Authors:** Adriane Guillaumin, Thibault Dhellemmes, Emma Perrot, Laura Boi, Daniel De Castro Medeiros, Christelle Glangetas, S. Dumas, Sandra Dovero, Nathalie Biendon, Elodie Ladeveze, Marc Landry, Erwan Bezard, Jérôme Baufreton, Gilberto Fisone, François Georges

**Affiliations:** Université de Bordeaux, CNRS, IMN, UMR5293 F-33000 Bordeaux, France; ORAMACELL, 75006 Paris, France; Department of Neuroscience, Karolinska Institutet, 17177 Stockholm, Sweden

**Keywords:** Dopaminergic neurons heterogeneity, bed nucleus of the stria terminalis, Amygdala, Vasoactive intestinal peptide, Central extended amygdala, Anxiety, NREM sleep, active-phase sleep stability, risk assessment, defensive behavior, Escape, immobility

## Abstract

Defensive behaviors are essential for survival, with risk assessment enabling organisms to evaluate and respond to potential threats. The dorsal raphe nucleus (DRN), a key neuromodulatory center, is crucial for encoding motivational salience and regulating arousal and sleep-wake states through its diverse neuronal populations, including dopaminergic neurons (DRN_DA_). While the roles of DRN_DA_ neurons have been studied, their specific contributions to threat evaluation are less understood. Recent research identifies a distinct subset of DRN_DA_ neurons that express vasoactive intestinal peptide (VIP) and project to the central amygdala (CeA) and the oval nucleus of the bed nucleus of the stria terminalis (ovBNST). Together, these two regions comprise the central extended amygdala, a key network regulating adaptive responses to threats. We hypothesized that distinct DRN_DA_ subpopulations exert diverging effects on sleep-wake regulation and that DRN_VIP_ neurons play a pivotal role in coordinating activity between the CeA and ovBNST, thereby influencing risk assessment and defensive response. To test this hypothesis, we used a combination of *in situ* hybridization, immunochemistry, whole-brain mapping, electrophysiology, and cell-specific genetic tools in mice and non-human primates. Our findings reveal that DRN_VIP_ neurons form a key DRN_DA_ neuronal subset, uniquely positioned to regulate the central extended amygdala through a feedback loop. These neurons receive inputs from Protein Kinase C delta (PKC-δ) neurons in the ovBNST and CeA and send glutamate-releasing projections back to these regions, modulating PKC-δ neuron excitability. Selective ablation of DRN_VIP_ neurons increases activity in both the BNST and CeA, disrupting active-phase sleep architecture and impairing risk assessment and defensive behaviors. Together, these findings suggest DRN_VIP_ neurons control specific phases of sleep and orchestrate the central extended amygdala’s role in risk assessment and defensive responses.

**HIGHLIGHTS:**

- DRN_VIP_ neurons form a subset of DRN_DA_ neurons in mice and non-human primates.
- DRN_VIP_ receive inputs from Protein Kinase C delta (PKC-δ) neurons in the ovBNST and CeA and project back to both.
- By releasing glutamate, DRN_VIP_ neurons regulate PKC-δ neuron excitability in the ovBNST and CeA.
- Ablating DRN_VIP_ neurons increases BNST and CeA activity, disrupts active-phase sleep architecture, and impairs threat responses.

**IN BRIEF:** DRN_VIP_ neurons, a key subset of DRN_DA_ neurons in mice and primates, are strategically positioned to influence the central extended amygdala via feedback loops. They regulate PKC-δ neuron excitability in the ovBNST and CeA through glutamate release, with their ablation heightening activity in these regions and altering active-phase sleep architecture, risk assessment and defensive behaviors.

## INTRODUCTION

Defensive behaviors are crucial for an organism’s survival in the face of threats. In response to perceived threats, mammals—and mice in particular— display a range of defensive behaviors, including freezing, fleeing, hiding, and defensive aggression (Blanchard et al., 1993). In a hostile or unfamiliar environment, risk assessment plays a central role in this repertoire of defensive responses (Blanchard and Blanchard, 1989). Characterized by cautious exploration and information gathering, this behavior allows the individual to evaluate potential threats and determine the appropriate response. The dorsal raphe nucleus (DRN), located within the ventromedial periaqueductal gray, is a major source of neuromodulators in the central nervous system (McDevitt et al., 2014) and plays a key role in regulating sleep homeostasis and diverse defensive behaviors, including aggression, escape responses, and possibly panic-like reactions (Miguel et al., 2010; Mobbs et al., 2007; Takahashi et al., 2022). It contains molecularly distinct neuronal subtypes, which are classified into four main groups: serotonergic (5-HT) neurons, followed by dopaminergic (DA), GABAergic, and glutamatergic neurons (Huang et al., 2019). Midbrain dopaminergic neurons are crucial for encoding responses to unexpected sensory events and processing aversive experiences (Horvitz, 2000; Schultz, 2010; Verharen et al., 2020), however, only a few studies have explored this question for dopaminergic neurons outside the ventral tegmental area (VTA) (Groessl et al., 2018), and none have specifically addressed their role in risk assessment and defensive behavior.

Significant progress in understanding the role of DA neurons in the brain has been made with the recognition of their heterogeneity (Roeper, 2013). This strategy over the past years has revealed that DA neurons are not a uniform population, but instead consist of subtypes with specialized roles, leading to a more nuanced comprehension of their contributions to neural circuits and behavior (Verharen et al., 2020). Over the past decades, research efforts have extensively focused on understanding the diversity of dopaminergic neurons in the VTA (de Jong et al., 2022), somewhat less so for those in the substantia nigra pars compacta (SNc) (Gaertner et al., 2022), and has only recently begun for dopaminergic neurons in the DRN (DRN_DA_ neurons)(Huang et al., 2019; Poulin et al., 2018).

Electrophysiological, molecular, and RNA sequencing approaches have revealed substantial heterogeneity within the DRN_DA_ neurons, encompassing diverse electrophysiological properties, gene expression profiles, as well as distinct morphological and spatial characteristics. However, recent studies exploring the function of DRN_DA_ neurons have largely overlooked their cellular heterogeneity, yet have revealed a surprisingly diverse range of roles for these neurons. These include roles in modulating social behavior (Matthews et al., 2016), mediating analgesia (Li et al., 2016), encoding aversive teaching signals (Groessl et al., 2018), controlling expression of incentive memory (Lin et al., 2020), promoting wakefulness (Cho et al., 2017; Lu et al., 2006), and regulating both positive and negative motivational salience (Cho et al., 2021; Cho et al., 2017), and more recently producing depressive phenotypes (Wang et al., 2024). This diversity suggests an equally varied DA neurocircuitry within the DRN. Moreover, these investigations primarily relied on two transgenic mouse lines used to specifically target and manipulate dopaminergic neurons that expressed Cre under control of dopamine transporter (DAT) or tyrosine hydroxylase (TH) genes. Although these lines are highly specific for DA neurons in the VTA and SNc, both lines exhibit limited specificity for DRN_DA_, with off-target effects on 30% to 50% of non-DA neurons (Cardozo Pinto et al., 2019).

Our primary goal in this study was to target a specific subpopulation of DRN_DA_ neurons using a transgenic mouse line with enhanced specificity. Recent studies have identified vasoactive intestinal peptide (VIP) mRNA, or VIP itself, as a molecular marker for a distinct subset of dopaminergic neurons within the DRN (Dougalis et al., 2012; Poulin et al., 2018; Zhao et al., 2022). Using intersectional genetic labeling strategies, Poulin et al. demonstrated that VIP+ DA neurons located in the DRN (DRN_VIP_ neurons) project to the central amygdala (CeA) and the oval nucleus of the bed nucleus of the stria terminalis (ovBNST) (Poulin et al., 2014). Studies in rodents, non-human primates, and humans have shown that the CeA and the ovBNST are key components of the central extended amygdala, essential for regulating adaptive responses to threats (Alheid and Heimer, 1988; Shackman and Fox, 2016). Both structures share a common cellular lineage, a striatal-like organization, and are predominantly composed of GABAergic neurons with similar structural and chemical features (Beyeler and Dabrowska, 2020). They also contain neurons expressing Protein Kinase C delta (PKC-δ), a marker critical for emotional processing and the modulation of defensive behaviors based on the proximity of a threat (Moscarello and Penzo, 2022; Williford et al., 2023). Early work by Davis and colleagues suggests that the anatomical connection between the ovBNST and CeA is the basis of their functional interaction (Davis et al., 1993; Lee and Davis, 1997). This functional interaction becomes particularly significant as a threat approaches, prompting a shift from a hypervigilant, anxiety-driven state governed by the BNST to a fear-driven state mediated by the amygdala (Avery et al., 2016; Shackman and Fox, 2016). However, rather than operating in a strictly sequential manner, recent findings suggest that the BNST and amygdala may be recruited simultaneously, indicating the involvement of a neural circuit capable of synchronizing the activity of both structures (Mobbs et al., 2020). These findings led us to hypothesize that DRN_VIP_ neurons form a homogeneous population that, due to their anatomical projections to both the ovBNST and CeA, occupy a strategic position capable of coordinating the activity of these regions, thereby controlling risk assessment and defensive behavior. Sleep stability modulates neural and cognitive processes essential for evaluating threats and selecting adaptive defensive behaviors (LaGoy et al., 2022). DRN_DA_ neurons have been characterized as a wake-promoting system (Cho et al., 2017; Leger et al., 2010; Lu et al., 2006). Optogenetic stimulation of these neurons rapidly awakens mice from sleep, while chemogenetic inhibition promotes non-REM (NREM) sleep even in the presence of arousing stimuli, suggesting that DRN_DA_ neurons serve as a "gatekeeper" for wakefulness during environmentally relevant events (Cho et al., 2017). Here, we investigate the function of DRN_VIP_ neurons in the context of sleep-wake states and threat-related behavior. In this context, we selected and validated the Vip-Cre transgenic mouse line to selectively target this specific subset of DRN_DA_ neurons projecting to the ovBNST and the CeA.

We used a combination of *in situ* hybridization and immunochemistry techniques in both mice and non-human primate tissues, along with comprehensive whole-brain mapping of cell type-specific inputs and outputs. Additionally, we use *in vivo* electrophysiology, patch-clamp methods for monitoring cell type-specific circuits, and cell type-specific genetic ablation to manipulate circuits. Our results show that, in both mice and primates, DRN_VIP_ neurons form an important subset of DRN_DA_ neurons. These neurons are strategically positioned to control the central extended amygdala through a feedback loop, receiving their main inputs from PKC-δ neurons in the ovBNST and CeA. They also send axonal collaterals to both the ovBNST and CeA, where they regulate PKC-δ neuron excitability via glutamate release. Selective genetic ablation of these neurons leads to increase *in vivo* electrophysiological activity in the BNST and CeA, and affects active-phase sleep architecture and threat responses. Taken together, we propose that DRN_VIP_ neurons represent a neural subset that coordinates activity within the central extended amygdala, controlling sleep stability and modulating risk assessment and defensive behaviors.

## RESULTS

### DRN_VIP_ neurons are a subset of DRN_DA_ neurons in mice and non-human primates

Here, we focused our study on the dopaminergic VIP-expressing neuronal population of the DRN. Using *in situ* hybridization, we demonstrated that approximately 50% of Th+ mRNA cells in the DRN also express VIP mRNA. Among these VIP+ neurons, the majority (83%) co-express Th+ in C57BL/6 mice (Supplementary Figure 1A). To further validate this approach, we employed the Vip-Cre mouse line to specifically target the DRN_VIP_ subpopulation, confirming their dopaminergic identity: 91% of the targeted cells expressed TH, and all (100%) co-localized with Cre recombinase (Supplementary Figure 1B). Interestingly, while *in situ* hybridization identified this dopaminergic phenotype, TH immunostaining of VIP-expressing neurons revealed a lower-than-expected expression of the TH enzyme (Figure 1A and B). This discrepancy suggests that VIP neurons exhibit low levels of TH protein, consistent with observations in other dopaminergic populations (Figure 1B; (Hokfelt et al., 1976)). Using Cre-dependent viral approaches in Vip-cre mice (Figure 1A) combined with the AdipoClear tissue-clearing technique, we selectively targeted DRN_VIP_ neurons and characterized the anatomical distribution of these neurons (Figure 1C-E). The overall organization of the DRN_VIP_ neuronal population resembles a "crab claw" with dense posterior and intermediate regions, accompanied by two extensions: a dorsal branch and a ventral branch, both oriented rostrally (Figure 1C-E). These DRN_VIP_ neurons are distributed along the lateral and ventral edges of the aqueduct of Sylvius, spanning the anteroposterior axis from -3.80 mm to -5.00 mm relative to bregma (Figure 1C-E). Quantification of GFP+ cell bodies in the DRN, performed on cleared brain samples (N=3) and analyzed with Imaris software revealed an average of 516 GFP+ neurons per brain (Figure 1F).

**Figure 1.**
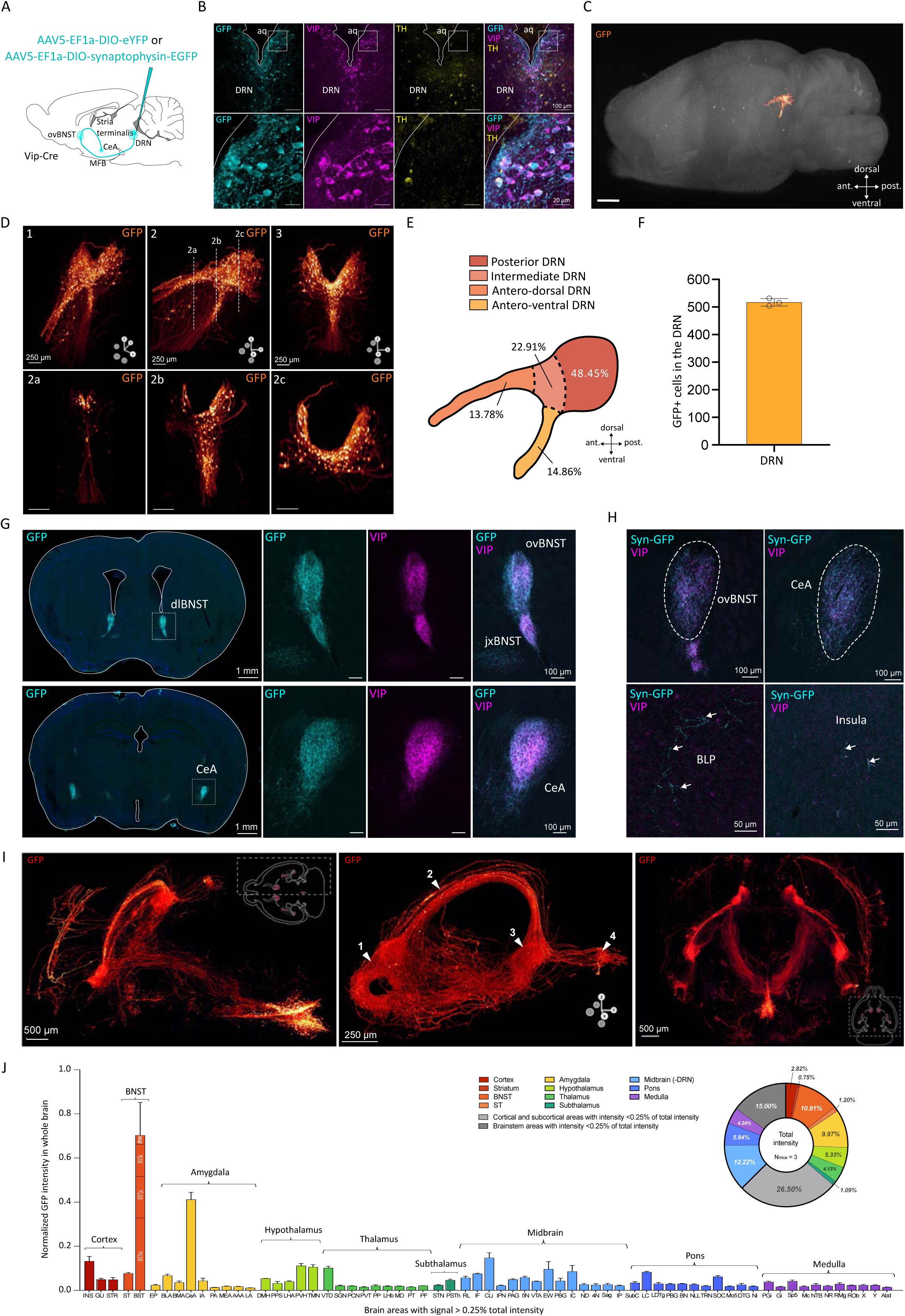
Anatomical characterization of DRN_VIP_ neurons and their projections. **(A)** Schematic representation of the injection site of the viral vector in Vip-Cre mice for labelling DRN_VIP_ neurons. **(B)** Confocal images of DRN_VIP_ neurons showing colocalization of GFP protein with VIP and in few cases with TH. **(C)** Reconstructed image of the DRN of a Vip-Cre mouse after viral transfection described in A and clearing technique. The image shows the location of DRN GFP positive cell bodies in the whole brain. **(D)** Images showing the distribution pattern of DRN GFP positive neurons from different view angles: frontal slightly angled (1), sagittal (2) and frontal (3). Images 2a, 2b an 2c corresponds to the three sections showed in image 2 at different antero-posterior levels from most anterior (2a) to most posterior (2c). **(E)** Schematic representing the distribution of GFP neurons in the DRN from a sagittal perspective and subdivided in different subparts: antero-dorsal, antero-ventral, intermediate and posterior. **(F)** Histogram showing the average number of transfected GFP positive cells in the DRN of Vip-Cre mice (N=3). **(G)** Epifluorescent images of GFP and VIP fibers in the dorsolateral BNST (dlBNST) and CeA from AAV5-EF1a-DIO-eYFP injection. **(H)** Confocal images of synaptophysin-EGFP expressed specifically in DRN_VIP_ neuron terminals in ovBNST, CeA, insula and BLP with VIP immunostaining. **(I)** Images of Adipoclear technique showing DRN_VIP_ cell bodies and projections through MFB and stria terminalis to BNST, CeA, BLP and insula from three different view angles, from left to right: a horizontal view of the right hemisphere, a sagittal view of the right hemisphere with white arrows pointing at the BNST (1), the stria terminalis (2), the CeA (3) and the BLP (4) and a horizontal view of both hemispheres. **(J)** Histogram showing a whole brain quantification (N=3) of GFP fiber intensities per mm^3^ with a threshold superior to 0.25% of total intensity associated to a donut representation including all brain areas without intensity threshold. Abbreviations: MFB: medial forebrain bundle; CeA: central nucleus of the amygdala; ovBNST: oval nucleus of the bed nucleus of the stria terminalis; jxBNST: juxtacapsular nucleus of the BNST; Syn-GFP: synaptophysin-GFP; BLP: posterior basolateral amygdala; VIP: vasoactive intestinal peptide. DRN: dorsal raphe nucleus; DA: dopaminergic; aq: aqueduct of Sylvius; TH: tyrosine hydroxylase. Allen Brain Atlas abbreviations were used for naming brain structures.

To investigate the existence of the DRN_VIP_ population in other species we performed VIP immunostaining on non-human primate coronal brain slices. DRN_VIP_ neurons of non-human primates exhibit a distribution similar to that observed in mice, aligning along the lateral and ventral regions of the aqueduct of Sylvius (Supplementary Figure 2A). Notably, double immunohistochemistry for TH and VIP in the DRN showed a high degree of co-localization between these markers, contrasting with the limited overlap observed in mice (Supplementary Figure 2B).

### DRN_VIP_ neurons are strategically positioned to influence the central extended amygdala via a feedback loop

DRN_DA_ neurons project to multiple regions, including the BNST, CeA, BLA, VTA, and hypothalamus (Cardozo Pinto et al., 2019; Matthews et al., 2016). Using AAV-EF1a-DIO-eYFP injections in the DRN (Figure 1A) combined with the AdipoClear tissue-clearing technique, we observed a highly restricted projection pattern for DRN_VIP_ neurons, with two primary output targets: the dorsolateral bed nucleus of the stria terminalis (dlBNST), including the oval (ovBNST) and juxtacapsular (juBNST) nuclei, and the central amygdala (CeA), particularly its lateral part (Figure 1G). Using an AAV-EF1a-DIO-synaptophysin-GFP virus injected into the DRN of Vip-Cre mice (Figure 1A and H), we confirmed that the observed labeling in the BNST, CeA, insular cortex, and posterior basolateral nucleus of the amygdala (BLP) corresponded to synaptic labeling rather than passing fibers. VIP immunostaining in NHPs revealed prominent VIP+ fibers in the ovBNST and CeA (Supplementary Figure 2C-F). In cleared mice brains (Figure 1, Supplementary Figure 3), GFP+ fibers leaved the DRN ventrally, joined the MFB, and reached the BNST where synaptic contacts were observed (Figure 1I). These fibers then traveled dorsally through the stria terminalis to innervate the CeA and, more caudally, the BLP. AdipoClear imaging also identified fibers diverging from the MFB before reaching the BNST, traveling directly to the CeA via the internal capsule. Secondary output targets included the BLP and insular cortex (Figure 1I). Quantitative fluorescence analysis confirmed strong projections to the dlBNST and anterior BNST, with the lateral CeA showing the highest fluorescence intensity among amygdala nuclei (Figure 1J). These results indicate that DRN_VIP_ neurons have a more restricted projection pattern compared to the entire DRN_DA_, with specific regions like the BLA excluded as targets. Images obtained from cleared brain samples strongly suggest that axons of DRN_VIP_ neurons sequentially innervate the BNST and then the CeA (Figure 1I). To confirm that individual DRN_VIP_ neurons can simultaneously project to both the BNST and CeA, we employed a modified rabies virus tracing strategy (Sun et al., 2023). Using a Cre-dependent helper virus lacking the glycoprotein (AAV1-EF1a-DIO-TVA950-T2A-WPRE) along with two rabies viruses injected into the ovBNST (pSADB19dG-GFP) and the CeA (PSADB19dG-mCherry) of Vip-Cre mice, we observed double-labeled cell bodies in the DRN (Supplementary Figure 4). This finding confirms that DRN_VIP_ neurons project to both the BNST and CeA (Supplementary Figure 4B).

### ovBNST_PKC-δ_ and CeA_PKC-δ_ are the main synaptic inputs to DRN_VIP_ neurons

To investigate the circuit in which DRN_VIP_ neurons are embedded, we employed a rabies-virus-based transsynaptic tracing approach in Vip-Cre mice to map their presynaptic inputs (Figure 2A). First, we confirmed the presence of starter cells co-expressing the helper virus and rabies reporter in the DRN (Supplementary Figure 5A-B). Surprisingly, our tracing experiment revealed that only two major brain structures provide substantial input to DRN_VIP_ neurons: the ovBNST and the lateral CeA (Figure 2B). While additional input arises from regions predominantly located in the hypothalamus and pons, these projections are comparatively sparse (Figure 2B). Previous studies have identified two main, non-overlapping neuronal populations within the ovBNST and CeA: one expressing protein kinase C delta (PKC-δ), which co-expresses D2 dopamine receptors, and another expressing corticotropin-releasing hormone (CRH) (Beyeler and Dabrowska, 2020; Wang et al., 2024). Since PKC-δ neurons constitute the largest subpopulation in both the ovBNST and CeA, we performed PKC- δ immunohistochemistry to characterize the identity of neurons projecting to DRN_VIP_ neurons. Our analysis revealed a strikingly high degree of colocalization between rabies-labeled neurons and PKC-δ expression, with 95.6% in the ovBNST and 95.4% in the CeA (Figure 2C-D). These findings reveal a highly specialized circuit in which DRN_VIP_ neurons receive dense input from two GABAergic structures, the ovBNST and CeA, while simultaneously projecting back to these same regions, forming a restricted feedback loop.

**Figure 2.**
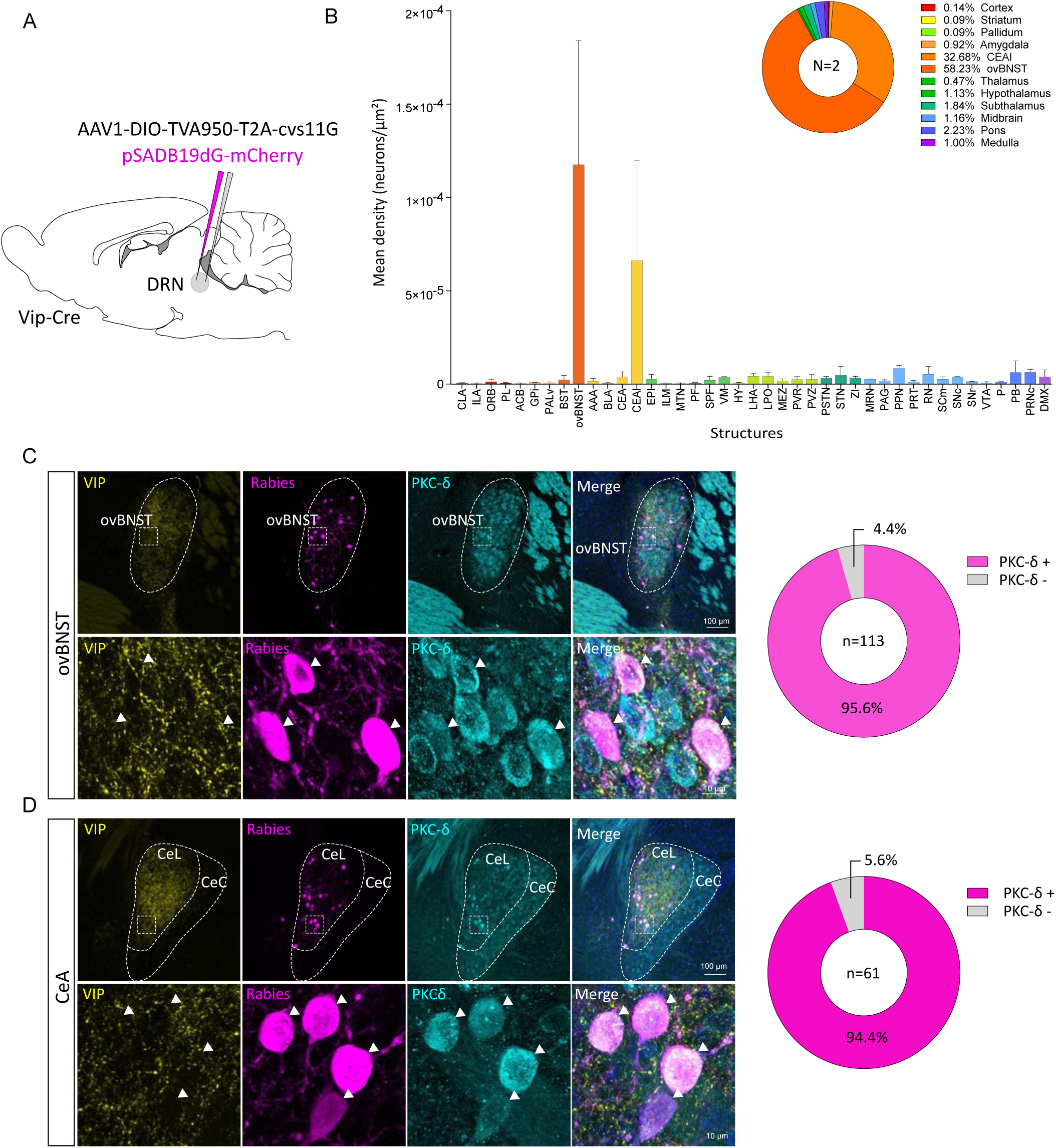
Characterization of DRN_VIP_ inputs. **(A)** Schematic representation of viral strategy to map inputs to DRN_VIP_ neurons using a Cre-dependant helper virus (AAV1-EF1a-DIO-TVA950-T2A-cvs11G) and a rabies virus (pSADB19dG-mCherry) in Vip-Cre mice. **(B)** Whole brain quantification of DRNVIP neurons inputs from rabies viral strategy. Only structures with a mean density superior to 5.10^-7^ are shown in the histogram. The pie chart shows mean densities of main brain regions. **(C)** Confocal images and donut graphs showing co-localization of the majority ovBNST and **(D)** CeA rabies positive neurons with PKC-δ protein (N=2 mice). Allen Brain Atlas abbreviations were used for naming brain structures.

### DRN_VIP_ neurons release glutamate

DRN_DA_ neurons have been shown to co-release dopamine and glutamate in downstream structures (Matthews et al., 2016). Here, we investigated whether DRN_VIP_ neurons also release glutamate. To address this, we used Vip-Cre/COP4 mice, in which channelrhodopsin is selectively expressed in VIP-expressing neurons. Brain slices containing the DRN, ovBNST, and CeA were prepared, and a blue-emitting optical fiber was placed either over DRN cell bodies to confirm depolarization upon light stimulation (Figure 3A) or over synaptic terminals in the ovBNST and CeA to assess neurotransmitter release (Figure 3D). We validated channelrhodopsin expression in DRN_VIP_ neurons by recording evoked inward currents upon light stimulation (Figure 3B-C). Optogenetic stimulation revealed that 44% of recorded neurons in the ovBNST (n = 11/25) responded, compared to only 6% in the CeA (n = 3/33) (Figure 3G). All responsive neurons exhibited excitatory post-synaptic currents (EPSCs), indicating a glutamatergic phenotype (Figure 3F). Furthermore, the addition of AMPA and NMDA receptor antagonists (CNQX and AP5) abolished these currents, confirming that DRN_VIP_ neurons release glutamate (Figure 3F). Characterization of synaptic transmission revealed low EPSC amplitudes and synaptic depression in both ovBNST and CeA neurons (Figure 3H). Given that a major neuronal population in these regions expresses PKC-δ, we examined whether DRN_VIP_ neurons preferentially target PKC-δ-expressing cells. Immunostaining for PKC-δ showed high colocalization with biocytin-filled neurons, reaching 82% in the ovBNST and 66% in the CeA, suggesting that DRN_VIP_ neurons predominantly innervate PKC-δ-expressing cells.

**Figure 3.**
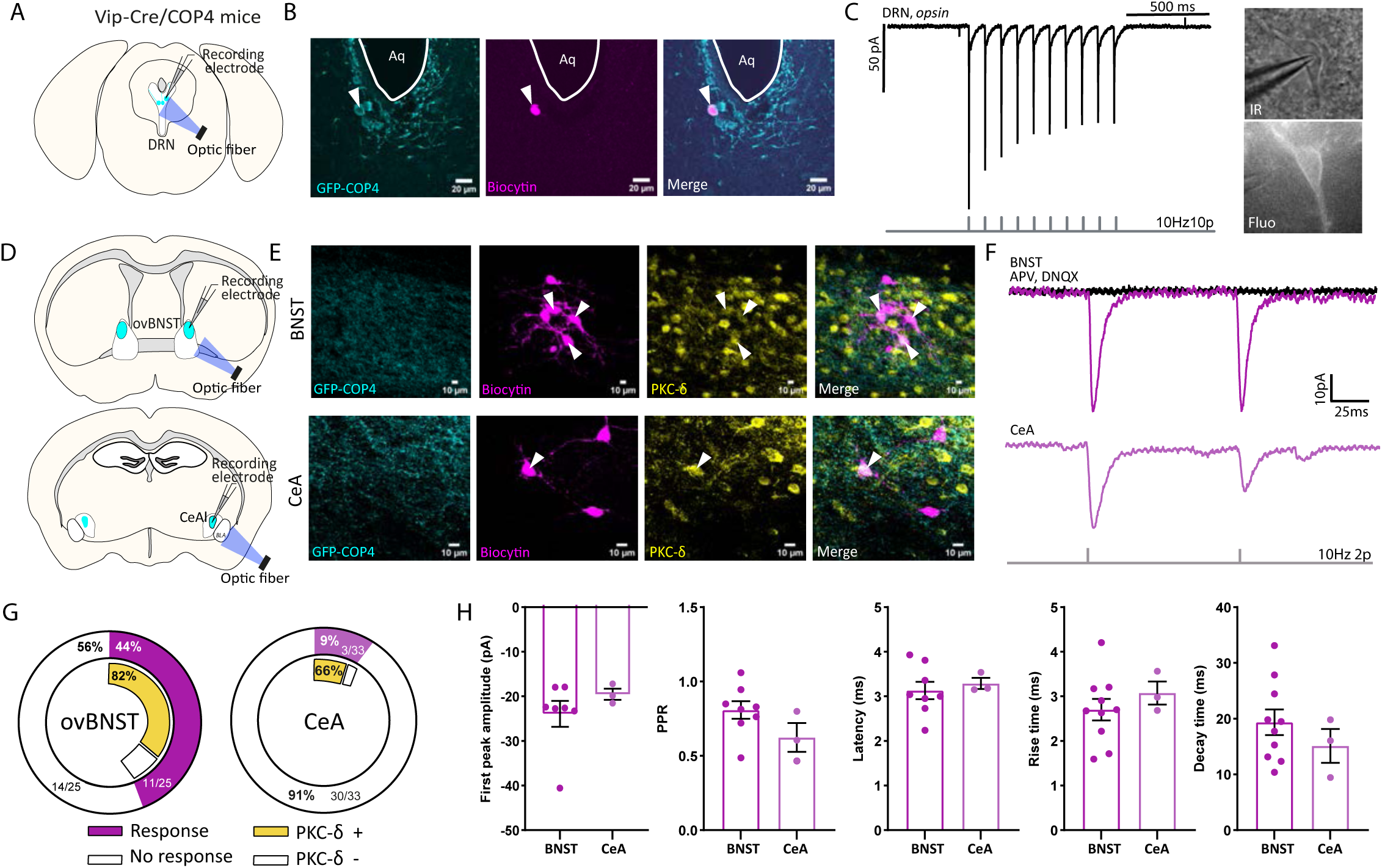
*Ex vivo* characterization of DRNVIP synaptic transmission to ovBNST and CeA. **(A)** Schematics of a DRN slice and positioning of recording electrode and optic fiber in Vip-Cre/COP4 mice. **(B)** Confocal images of a patched GFP+ neuron filled with biocytin in the DRN of a Vip-Cre/COP4 mouse **(C)** Channelrhodopsin evoked currents in a DRN GFP+ neuron associated to fluorescent and infra-red images **(D)** Schematics of ovBNST and CeA slices and positioning of recording electrode and optic fiber in Vip-Cre/COP4 mice. **(E)** Confocal images of patched neurons filled with biocytin and PKC-δ immunostaining in ovBNST and CeA. **(F)** Excitatory post-synaptic evoked currents recorded in ovBNST and CeA neurons upon optogenetic stimulation of DRN_VIP_ fibers. Purple curves show inward currents abolished by application NMDA antagonists, APV and CNQX (black curve). **(G)** Donut graphs showing the proportion of neurons responding to optogenetic stimulation of DRN_VIP_ fibers colocalizing with PKC-δ. **(H)** *Ex vivo* electrophysiological characterization of ovBNST and CeA excitatory post-synaptic currents evoked by optogenetic stimulation of DRN_VIP_ fibers. Abbreviations: DRN: dorsal raphe nucleus; ovBNST oval nucleus of the bed nucleus of the stria terminalis; Aq: aqueduct of Sylvius; CeA: central nucleus of the amygdala; PPR: paired-pulse ratio.

### DRN_VIP_ neuron ablation leads to sleep stability

Population activity of DRN_DA_ neurons is closely correlated with sleep-wake states, and chemogenetic inhibition of these neurons reduces responsiveness to external stimuli, impairing the ability to awaken from sleep (Cho et al., 2017). Moreover, chemical lesions targeting these cells result in the onset of severe hypersomnia (Lu et al., 2006). Building on these findings, we investigated whether ablation of DRN_VIP_ neurons could disrupt sleep architecture by altering the balance between wakefulness and sleep states. Specifically, we hypothesized that modulating the activity of these neurons could impact the stability and duration of NREM episodes, ultimately influencing the overall sleep-wake cycle. To determine whether DRN_VIP_ dopamine subgroups exert distinct effects on sleep-wake regulation (Cho et al., 2017), we selectively ablated DRN_VIP_ neurons using an AAV5-EF1a-FLEX-taCaspase3 viral construct (Casp3), while control mice received an AAV5-EF1-DIO-eYFP virus (Ctrl) in the DRN of Vip-Cre mice (Figure 4A). Histological analysis, conducted after a series of behavioral tests (Figure 4B), confirmed that Casp3 ablation effectively eliminated VIP**+** neurons in the DRN (Figure 4C). Additionally, VIP and TH fiber density was markedly reduced in the ovBNST and CeA (Figure 5D-E), further supporting the dopaminergic phenotype of DRN_VIP_ neurons despite their low TH expression in cell bodies.

**Figure 4.**
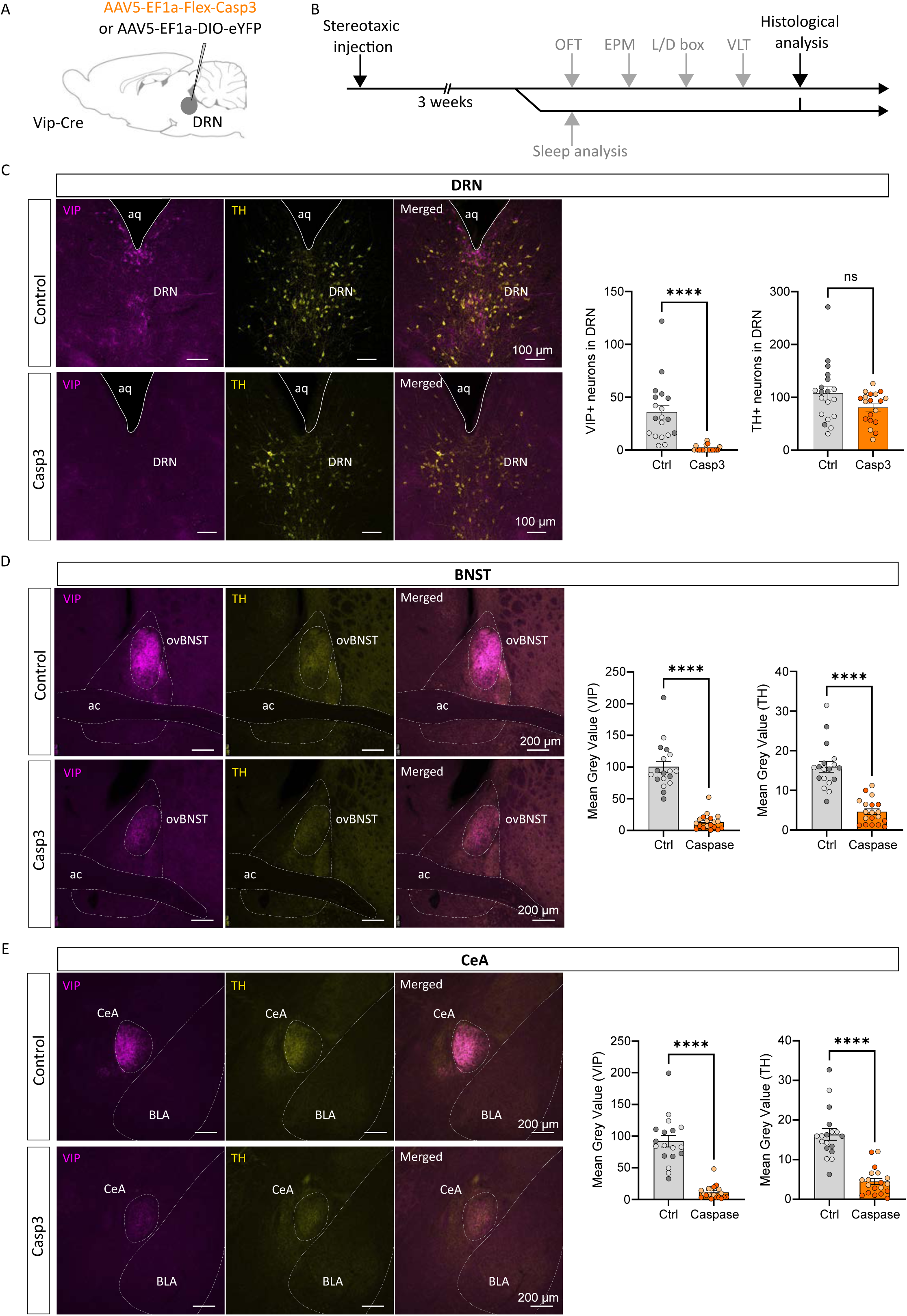
Histological analysis of behavioral experimental group. **(A)** Schematic representation of AAV5-EF1-Flex-Casp3 injection in the DRN of Vip-Cre mice and control virus. **(B)** Chronological timeline of the experimental procedure from virus injection to histological analysis. Mice were separated in two batches, one for assessing anxiety-related behaviors and defensive behaviors and one for sleep analysis. **(C)** Confocal images and quantification of TH+ and VIP+ neurons in the DRN of AAV5-EF1-Flex-Casp3 injected mice (referred as Casp3) and AAV5-EF1a-DIO-eYFP injected mice (referred as Ctrl) (Mann-Whitney test, p-value<0.0001 for VIP+ quantification; Unpaired T-test, p-value=0.0753 for TH+ quantification). **(D)** Confocal images and quantification of mean intensity of TH+ and VIP+ fibers in the ovBNST of Casp3 mice and controls (Mann-Whitney test, p-value<0.0001 for both VIP+ and TH+ quantification). **(E)** Confocal images and quantification of mean intensity of TH+ and VIP+ fibers in the CeA of Casp3 mice and controls (Mann-Whitney test, p-value<0.0001 for both VIP+ and TH+ quantification). Graphs show mean ± SEM. Abbreviations: OFT: open field test; EPM: elevated plus maze; L/D box: light/dark box; VLT: visual looming test; DRN: dorsal raphe nucleus; ovBNST: oval nucleus of the bed nucleus of the stria terminalis; CeA: central nucleus of the amygdala; BLA: basolateral nucleus of the amygdala; aq: aqueduct of Sylvius; ac: anterior commissure.

**Figure 5.**
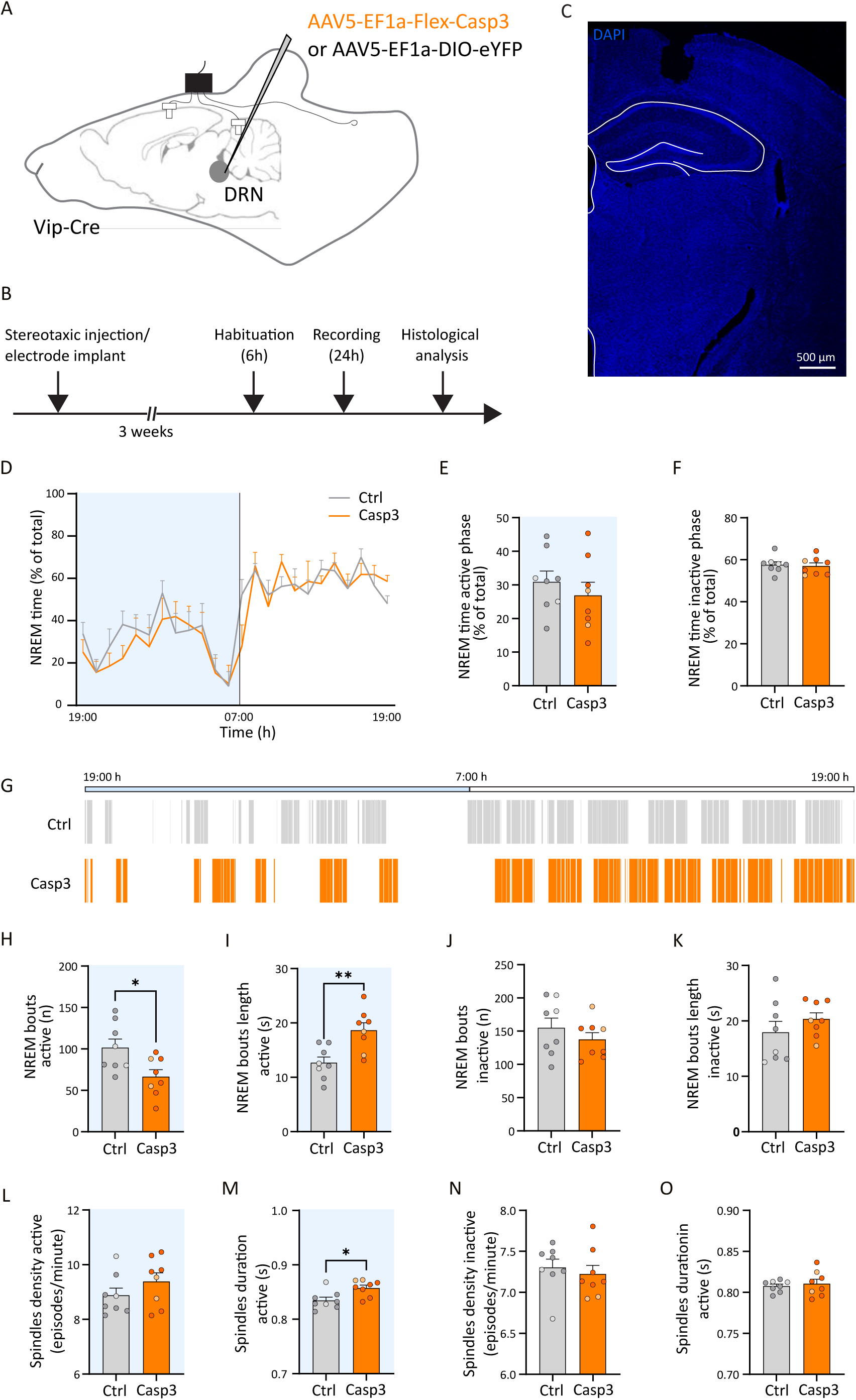
Effect of DRN_VIP_ genetic ablation on NREM state in Vip-Cre mice. **(A)** Schematic representation showing AAV5-EF1a-Flex-Casp3/AAV5-EF1a-DIO-eYFP injections in the DRN of Vip-Cre mice, and EEG, reference and EMG placement. **(B)** Experimental timeline of sleep analysis. **(C)** Representative confocal picture showing cortical EEG positioning. **(D)** Graphics showing NREM sleep percent time during active (19- to 7 h) and inactive (7- to 19 h) phases in Ctrl and Casp3 groups. **(E, F)** Bar graphs showing NREM sleep percent during the active (19- to 7 h), (E, Unpaired T-test p-value=0.4375) and the inactive (7- to 19 h), (F, Unpaired T-test p-value=0.7605) phases in Ctrl and Casp3 groups. **(G)** Representative illustration of the NREM sleep bouts distribution in Ctrl and Casp3 mice during the 24-hour circadian cycle. Bars indicate NREM sleep episode occurrences. **(H, I)** Bar graphs showing the number (H, Unpaired T-test p-value=0.0207) and the length (I, Unpaired T-test p-value=0.0038) of NREM sleep bouts during the active phase (19- to 7 h) in Ctrl and Casp3 groups. **(J, K)** Bar graphs showing the number (J, Unpaired T-test p-value=0.3486) and the length (K, Unpaired T-test p-value=0.3050) of NREM sleep bouts during the inactive phase (7- to 19 h) in Ctrl and Casp3 groups. **(L, M)** Bar graphs showing NREM sleep spindles density (L, Unpaired T-test p-value=0.2357) and duration (M, Unpaired T-test p-value=0.0118) during the active phase (19- to 7 h) in Ctrl and Casp3 groups. **(N, O)** Bar graphs showing NREM sleep spindles density (N, Unpaired T-test p-value=0.5951) and duration (O, Unpaired T-test p-value=0.6303) during the inactive phase (7- to 19 h) in Ctrl and Casp3 groups. Light circles in bars correspond to female mice while dark circles correspond to male mice. *p-value < 0.5, **p < 0.01. Graphs show mean ± SEM.

Caspase-induced ablation of DRNVIP neurons resulted in significant alterations of sleep architecture occurring during the active phase of the 24-hour recording period (Figure 5). Specifically, the number of NREM sleep episodes decreased, while their average duration increased (Figure 5H-I), resulting in enhanced sleep stability in comparison to control. This increase in stability occurred in concomitance with changes in NREM spindles (Dang-Vu et al., 2010; Kam et al., 2023), which are brief 10-15 Hz oscillatory events implicated in the modulation of sensory inputs and spontaneous sleep disruptions (Fernandez and Luthi, 2020). We observed a significant increase in spindle duration (Figure 5M), along with evidence suggesting a contribution of spindle density to NREM sleep regulation (Figure 8A’). Importantly, total NREM sleep time remained unchanged (Figure 5J-K), excluding the occurrence of excessive daytime sleepiness (EDS). In line with the decreased number of NREM episodes, we observed reduced number of bouts in the AWAKE state (Supplementary Figure 6D). No differences were observed in the duration of wakefulness or REM sleep (Supplementary Figure 6).

### DRN_VIP_ neuron ablation changes risk assessment in anxiety-related tests

Both human and rodent studies have demonstrated that the BNST and CeA play a key role in threat monitoring, including anxiety-like behaviors, risk assessment, and defensive responses (Avery et al., 2016; Groessl et al., 2018; Mobbs et al., 2020). Given our finding that the ovBNST and CeA (lateral division) are the two main synaptic targets of DRN_VIP_ neurons, we employed the same genetic ablation approach (Figure 4) to investigate their role in anxiety, risk assessment, and defensive behaviors. To assess how the loss of these neurons affected BNST and CeA activity, we performed *in vivo* single-cell electrophysiological recordings. Genetic ablation of DRN_VIP_ neurons led to a significant increase in neuronal firing frequency in both regions (Supplementary Figure 7). To examine the behavioral consequences of DRN_VIP_ neuron ablation, we assessed anxiety-related and risk assessment behaviors using the open field test (OFT), elevated plus maze (EPM), and light/dark box (L/D box) (Figure 6A-B). No differences were observed between Casp3 and control mice in the OFT (Figure 6C). In the EPM, Casp3 mice spent significantly less time in the open arms (OA) and exhibited more protected head dips, indicating increased risk assessment behavior (Figure 6D) (Rodgers and Dalvi, 1997; Rodgers and Johnson, 1995). However, no differences were found in other risk assessment or anxiety-related measures, such as stretch-attend postures (SAP), center time, closed-arm entries, or total distance traveled (Figure 6D). In the L/D box test, Casp3 mice did not show reduced time in the lighted compartment, and none of the measured variables significantly differed from controls (Figure 6E).

**Figure 6.**
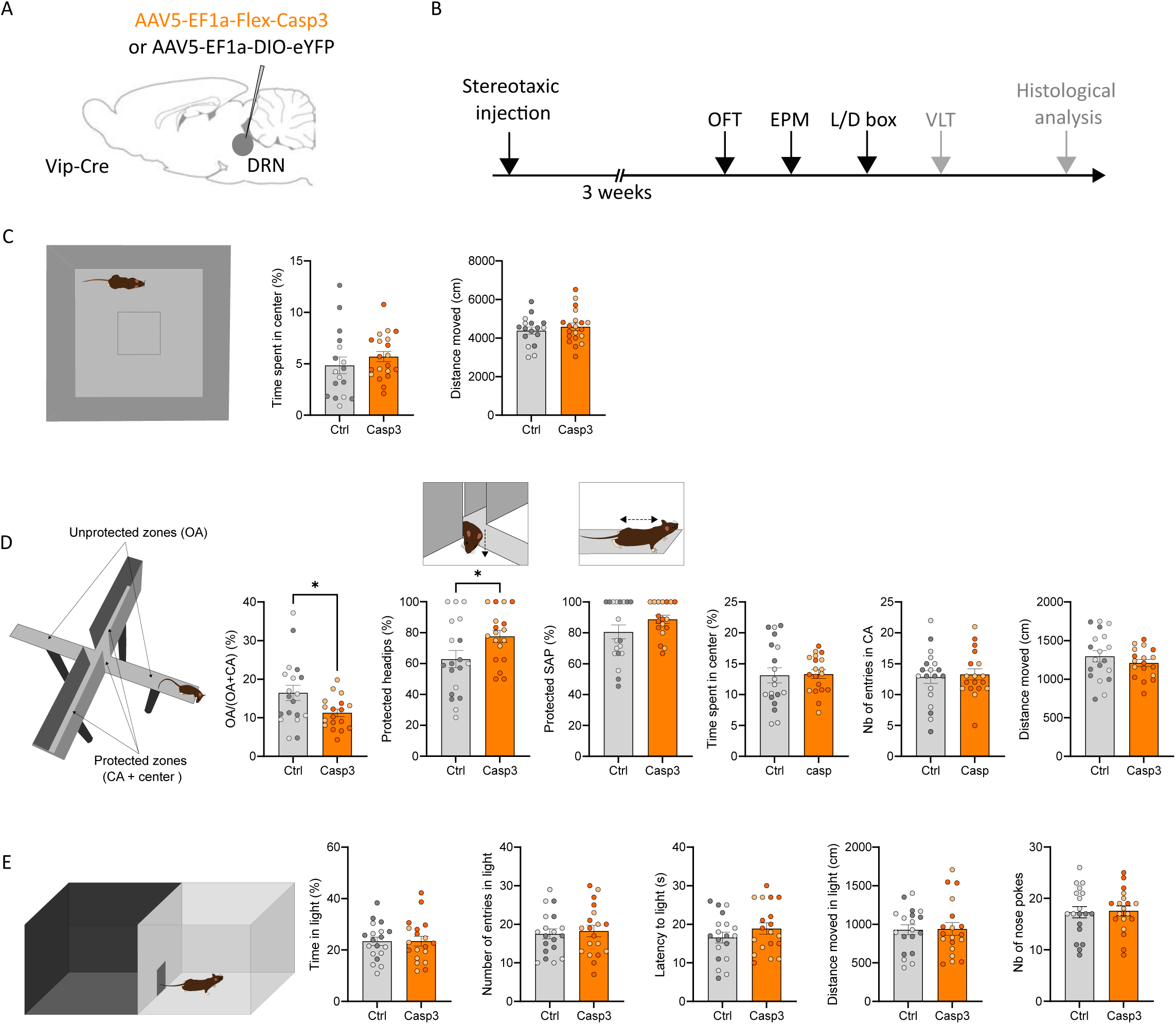
Effect of DRN_VIP_ genetic ablation on risk assessment, anxiety and locomotion in Vip-Cre mice. **(A)** Schematic representation of AAV5-EF1a-Flex-Casp3 or control viruses in the DRN in Vip-Cre mice. **(B)** Chronological timeline of the experimental procedure from virus injection to the three behavioral tests depicted below. **(C)** Schematic representation of the Open Field Test (OFT) arena. Quantification of the time in spent in center (Unpaired T-test, p-value=0.3591) and distance moved (Unpaired T-test, p-value=0.4645) between control and Casp3 mice. **(D)** Schematic representation of the Elevated Plus Maze (EPM) arena with arrows indicating protected (closed arms CA and center) and unprotected (open arms OA) zones. Quantification of the time spent in OA expressed in ratio (Unpaired T-test p-value=0.0267), protected headips (Unpaired T-test p-value=0.0396), protected SAP (Mann-Whitney test, p-value=0.2841), time spent in center (Unpaired T-test p-value=0.8845), number of entries in CA (Unpaired T-test p-value=0.7502) and distance moved (Unpaired T-test p-value=0.329) between control and Casp3 mice. **(E)** Schematic representation of the Light/Dark box (L/D box) arena. Quantification of the time spent in the light compartment (Unpaired T-test p-value=0.9668), number of entries in light compartment (Unpaired T-test p-value=0.7103), latency to light compartment (Unpaired T-test p-value=0.2553), distance moved in light compartment (Unpaired T-test p-value=0.9202) and number if nose pokes (Unpaired T-test p-value=0.8581). Light circles in graphs correspond to female mice while dark circles correspond to male mice. *p-value < 0.5. Graphs show mean ± SEM.

### DRN**_VIP_** neurons are required for adaptive escape responses to looming threats

The BNST is critical for threat anticipation, particularly in response to uncertain threats (Grogans et al., 2024; Hulsman et al., 2024; Hulsman et al., 2021; Mobbs et al., 2010; Siminski et al., 2021). To assess this function in Casp3 mice, we employed a visual looming test (VLT) (Figure 7A-C) (Daviu et al., 2020). In this task, a visual stimulus mimicking an approaching predator is triggered when the mouse enters a designated zone, prompting escape or other defensive behaviors (Figure 7B). The looming stimulus was specifically designed to favor escape responses over freezing. Casp3 mice exhibited a significantly lower probability of escape compared to controls across all three testing days (Figure 7D). Control mice displayed escape responses in 74% of trials, whereas Casp3 mice escaped in only 40% of trials (Figure 7E). Furthermore, escape vigor—quantified as the velocity of the return to the shelter—was reduced in Casp3 mice compared to controls. While control mice returned to the shelter with increased velocity, Casp3 mice moved at a similar speed during both the outward trip and the escape (Figure 7F). Escape latency at the first escape trial was similar between groups and decreased over repeated testing (Figure 7G). In addition to reduced escape probability, Casp3 mice spent less time in the shelter and more time in the trigger zone than control mice (Figure 7H). Since both control and Casp3 mice were exposed to the visual cue between 7 and 10 times on Day 1 of the VLT, we hypothesized that the VLT chamber had acquired an aversive valence by Day 2. However, no significant differences were observed between groups in the time spent in the trigger zone during the habituation phase on the second day (H2) of the VLT (Figure 7I). To further investigate risk assessment, we quantified rearing behavior, a relevant measure in the VLT since the visual stimulation mimics an overhead airborne predator. No differences in rearing behavior were observed between groups during habituation of day 1 (H1) (Figure 7J); however, Casp3 mice performed significantly more rearing both before and after visual cues, suggesting increased risk assessment (Figure 7J). To assess defensive responses, we measured cumulative immobility within 7s of stimulus onset across 10 trials. Casp3 mice spent significantly less time immobile than controls (Figure 7K). Importantly, no differences were observed in general locomotor activity between groups (Figure 7L).

**Figure 7.**
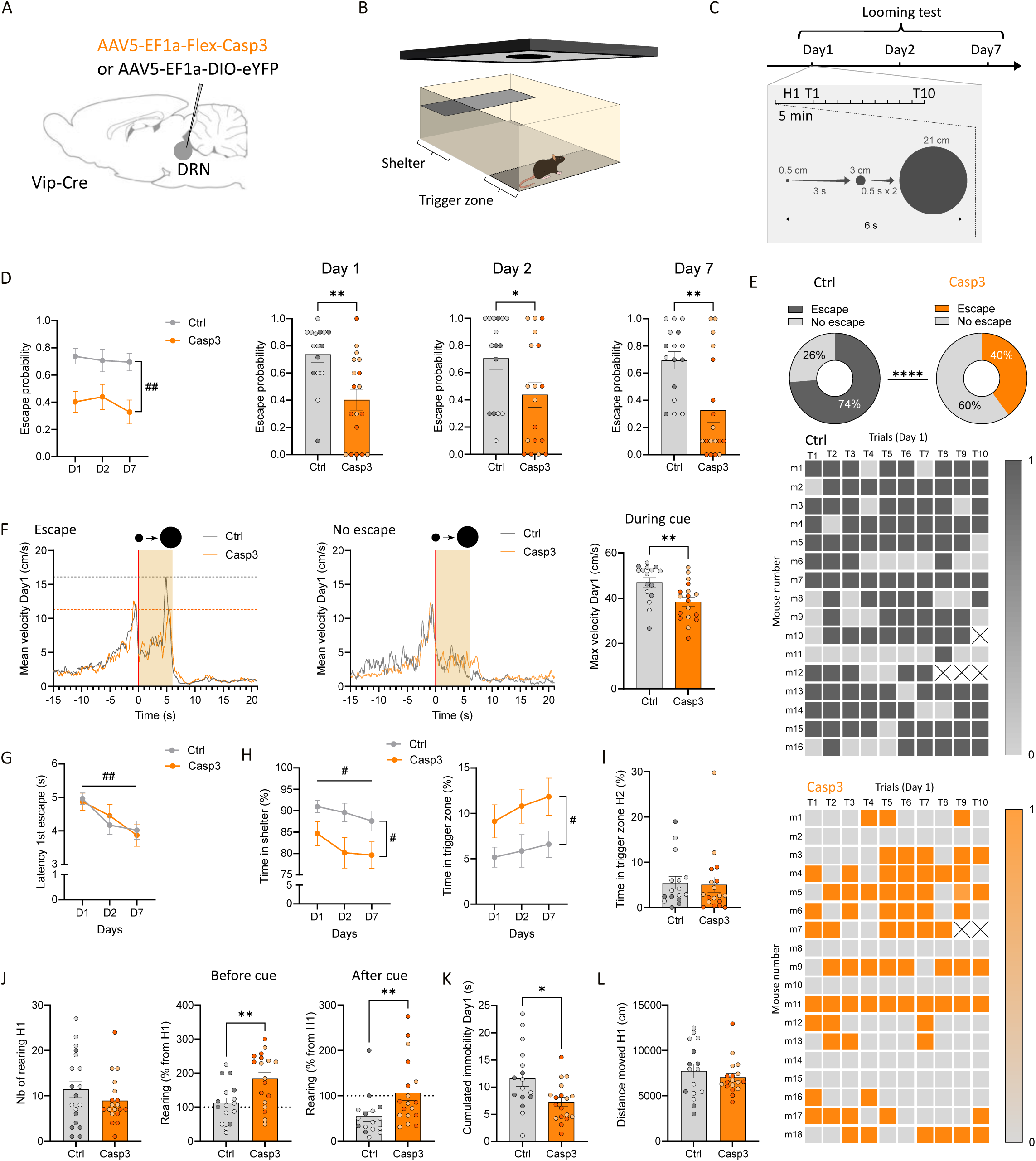
Effect of DRN_VIP_ genetic ablation on defensive behaviors in Vip-Cre mice. **(A)** Schematic representation of AAV5-EF1a-Flex-Casp3 and control virus injections in the DRN in Vip-Cre mice. **(B)** Schematic representation of the visual looming test setup. **(C)** Experimental timeline of the visual looming test with a detailed view of the procedure for Day 1 showing 5 min of habituation followed by 10 cues (=10 trials) represented by a black expanding round. **(D)** Quantification of the escape probability between control and Casp3 mice along days (Mixed-effects analysis, F_group_ (1, 32) = 10.20, p-value=0.0031) and for each day (Mann-Whitney test, Day1 p-value=0.0013, Day2 p-value=0.0423, Day7 p-value=0.0034). **(E)** On the top, donut graphs showing the percentage of escaping and not escaping trials in control mice (left) and Casp3 mice (right) (Fisher’s exact test, p-value<0.0001). Below, two heatmaps showing the detailed behavior at Day 1, escaping (dark color) or not escaping (light color) for each mouse and for each trial with control mice in the top heatmap and Casp3 mice in the bottom heatmap. Trials with a cross mean that the mouse did not perform the trial. **(F)** Graphs showing the average velocity of control and Casp3 mice at Day 1 for “escape” trials on the left and no escape trials on the right. The histogram on the right shows the max velocity at Day 1 for “escape” trials, representing the escape vigor (Mann-Whitney test, p-value=0.0056). In these three graphs only, the velocity threshold for “escape” trials was set on 20 cm/s to measure the vigor of the return to the shelter. No escape trials correspond to a velocity below 20 cm/s for the return to the shelter or the absence of return to the shelter. Light orange rectangles correspond to the period when the visual looming cue is presented. **(G)** Graph showing the latency of the first escape at Days 1, 2 and 7 (Mixed-effects analysis, F_Days_ (1.928, 53,03) = 7.372, p-value=0.0017). **(H)** Graphs showing the total time in shelter (2Way RM ANOVA, F_Group_ (1, 32) = 5.343 p-value=0.0274, F_Days_ (1.892, 60,53) = 3.747 p-value=0.0314) and trigger zone (2Way RM ANOVA, F_Group_ (1, 32) = 4.616, p-value=0.0393) at Days 1, 2 and 7. **(I)** Graph showing the time in trigger zone during the 5 min habituation of Day 2 (H2) (Mann-Whitney test, p-value=0.4174). **(J)** Graphs showing quantification of rearing behavior during the 5 min habituation of Day 1 (H1) (Mann-Whitney test, p-value=0.4375) and as a percentage from H1 before and after the visual looming cue (Before cue: Unpaired T-test, p-value=0.0062, One sample T-test, p-value=0.3974 for controls and 0.0003 for Casp3; After cue: Mann-Whitney test, p-value=0.0046, One sample T-test, p-value=0.0013 for controls and 0.6910 for Casp3). **(K)** Graph showing the cumulated immobility at Day 1 during the visual looming cues (Unpaired T-test, p-value=0.0135). **(L)** Graph showing the distance moved during the 5 min habituation of Day 1 (H1) (Mann-Whitney test, p-value=0.6212). Light circles in graphs correspond to female mice while dark circles correspond to male mice. Significant effects using 2Way RM ANOVA or Mixed-effects analyses are represented with a hash symbol. Significant effects using Unpaired t-test or Mann-Whitney test are represented with an asterisk symbol. #p-value < 0.05, ##p-value < 0.01, *p-value < 0.5, **p-value < 0.01.

### DRN_VIP_ neurons are essential for controlling sleep stability and coordinating risk assessment and defensive responses

To systematically evaluate behavioral changes following DRN_VIP_ neuron ablation, we computed z-scores across multiple tests, categorizing behaviors into sleep stability, defensive behavior, risk assessment, anxiety and locomotion (Figure 8) (Guilloux et al., 2011). Interestingly, Casp3 mice exhibited a significant increase in NREM sleep specifically during the active phase (Figure 8A and 8A’), but not the inactive phase (Figure 8B and 8B’). Additionally, they showed a reduction in defensive behaviors together with an increase in risk assessment (Figure 8C–D and 8C’–D’). However, no significant differences were found in anxiety-related behaviors, despite the reduced open-arm time in the EPM or locomotion (Figure 8E-F and 8E’-F’). Together, these findings suggest that DRN_VIP_ neurons play a crucial role in controlling sleep stability and balancing threat assessment and defensive responses, ensuring an adaptive response to visual threats.

**Figure 8.**
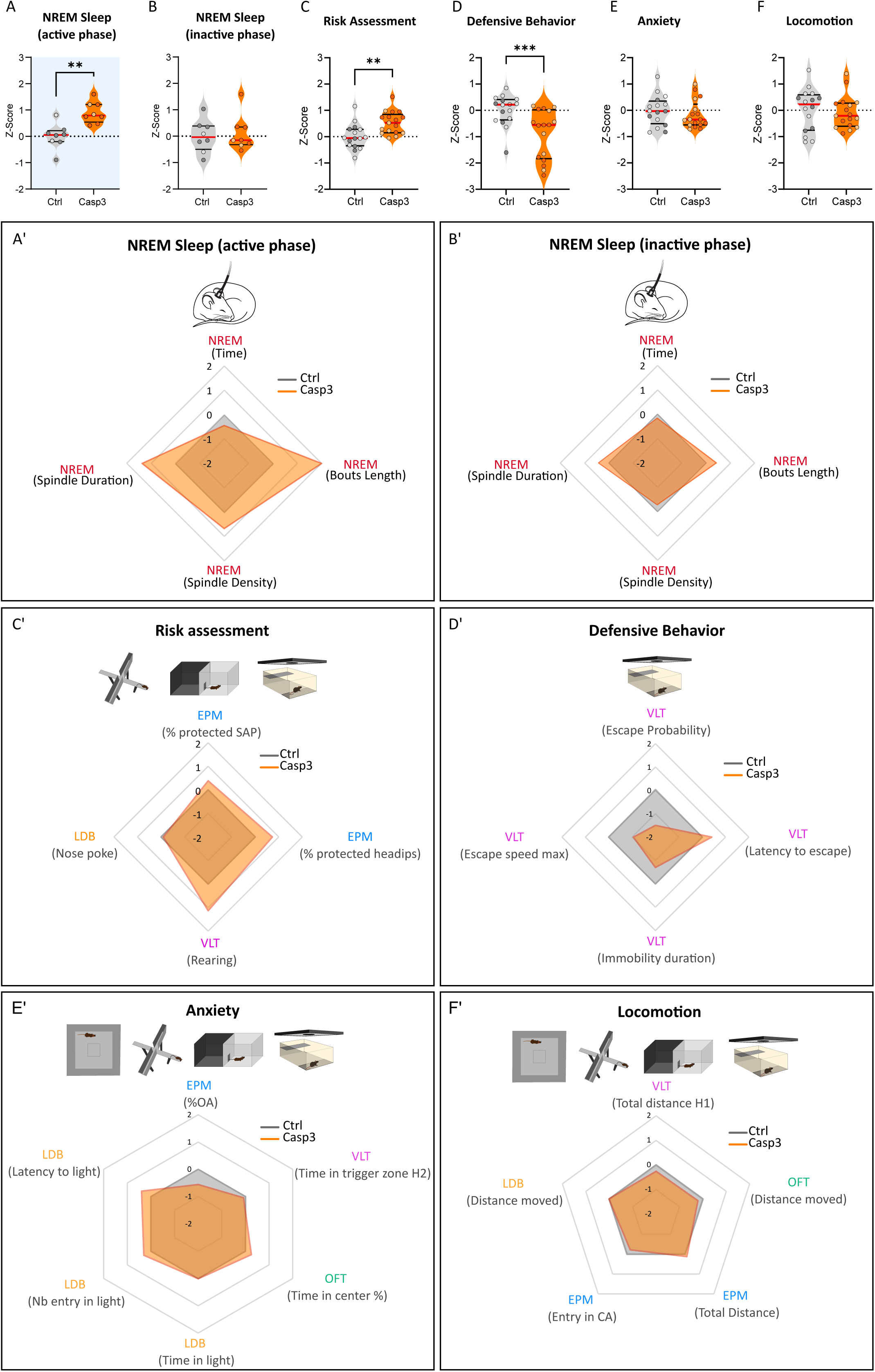
Scoring of the effect of DRN_VIP_ genetic ablation on behavior. Graph showing Z-score of NREM sleep (active phase (**A**) and inactive phase (**B**)), risk assessment (**C**), Defensive Behavior (**D**), Anxiety (**E**) and Locomotion (**F**) measured from several variables from different behavioral tests as shown in **A’** for NREM sleep (active phase) and **B’** for NREM sleep (inactive phase), **C’** for risk assessment, **D’** for defensive behavior, **E’** for anxiety and **F’** for locomotion (NREM sleep (active phase), p-value=0.012; NREM sleep (inactive phase), T-test, p-value=0.7542), Risk assessment, Unpaired T-test, p-value=0.0075; Defensive behavior, anxiety and locomotion, Mann-Whitney, respective p-values=0.009, 0.4654 and 0.5334). Selected variables for each test are shown in parenthesis below the test. Abbreviations: EPM: elevated plus maze; LDB: Light/Dark box; VLT: visual looming test; OFT: open field test; SAP: stretch attend posture; pHD: protected headips; H1: Habituation Day1; H2: Habituation Day2; OA: open arms; CA: closed arms.

## DISCUSSION

Evaluating risks in unfamiliar, complex, or hazardous environments and selecting appropriate behavioral responses to mitigate potential threats are essential for animal survival and may be influenced by homeostatic sleep pressure (Loftus et al., 2022). In this study, we present a comprehensive analysis of a relatively unexplored subpopulation of DA neurons located in the midbrain. Our findings show that, in both mice and primates, VIP- expressing neurons represent a significant subset of DA neurons within the region encompassing the ventral periaqueductal gray and dorsal raphe. In mice, this population of approximately 500 DRN_VIP_ neurons, representing 40% of all DRN_DA_ neurons, is located in a crab-claw-shaped region beneath the cerebral aqueduct, occupying a volume of 0.155 mm³. This specific neuronal population is strategically positioned to regulate the central extended amygdala through a feedback circuit. It primarily receives input from PKC-δ neurons in ovBNST and the CeA (lateral division). In turn, DRN_VIP_ neurons send axonal projections back to both the ovBNST and CeA (lateral division), where they modulate PKC-δ neuronal activity via glutamate release. Selective genetic ablation of DRN_VIP_ neurons leads to several key effects: 1) increased electrophysiological activity in the BNST and CeA, 2) disruption of sleep architecture during the active phase, characterized by a decrease in the frequency of NREM episodes and an increase in their duration, which suggests enhanced sleep stability without a rise in total NREM sleep time or excessive daytime sleepiness, and 3) impairment of risk-assessment and defensive responses. These findings offer valuable insight into the neural circuits controlling sleep stability and underlying risk-assessment behavior under normal physiological conditions.

### DRN_VIP_ neurons are strategically positioned to influence the central extended amygdala via a feedback loop

Our study revealed that DRN_VIP_ neurons are a critical subset of DRN_DA_ neurons in mice and non-human primates. It has been found that DRN_VIP_ neurons primarily project to the CeA and BNST (Poulin et al., 2018; Zhao et al., 2022). Our findings further reveal that the dorsolateral BNST (including the ovBNST and juxtacapsular BNST), anterior BNST, and the CeA (lateral division) are the principal target regions of DRN_VIP_ neurons, followed by the insular cortex and posterior basolateral amygdala. Our *in situ* hybridization results indicate that DRN_VIP_ neurons can be classified as a subset of DRN_DA_ neurons, given their colocalization with Th mRNA in over 80% of cases. Based on their molecular profile, three distinct subsets of dopaminergic neurons have been identified within the dorsal DRN (Huang et al., 2019). Our findings further confirm this heterogeneity and reveal that DRN_VIP_ neurons constitute 40% of the total dopaminergic neuron population in the DRN. The classification of DRN_VIP_ neurons as dopaminergic has been a topic of recent debate (Zhao et al., 2022). Our observations in mice reported that these neurons exhibit weak immunohistochemical detection of TH (Figure 1B). However, this does not appear to be the case in non-human primates, where DRN_VIP_ neurons show high levels of TH expression (Supplementary figure 2). Although there is a clear link between TH expression levels and the ability of dopamine neurons to synthesize and release dopamine, this relationship is complex and influenced by regulatory mechanisms as well as compartmental differences within neurons (Salvatore et al., 2016). A more targeted investigation will be required to conclusively determine whether DRN_VIP_ neurons modulate the activity of ovBNST and CeA (lateral division) neurons via dopamine release. Both the ovBNST and CeA (lateral division) are predominantly composed of GABAergic neurons, yet they are both highly heterogeneous in terms of cellular composition. For instance, PKC-δ-expressing neurons represent a distinct subpopulation with a similar organizational pattern in both the ovBNST and CeA (lateral division) (Ye and Veinante, 2019). Our findings reveal that DRN_VIP_ neurons are synaptically connected to PKC-δ neurons and use glutamate as a neurotransmitter in both the ovBNST and CeA (Figure 4). Moreover, over 90% of the synaptic inputs to DRN_VIP_ neurons originate from the ovBNST and CeL, and approximately 95% in the ovBNST and in the CeA (lateral division) of the neurons synaptically connected to DRN_VIP_ neurons are PKC-δ-expressing cells (Figure 3). PKC-δ neurons in the BNST and CeA (lateral division) are proposed to constitute a parallel microcircuit within the central extended amygdala (Cai et al., 2014; Ye and Veinante, 2019). BNST PKC-δ neurons are activated during risk assessment (Williford et al., 2023) and, together with CeA (lateral division) PKC-δ neurons, may form a retrograde inhibitory pathway that acts as a brake to prevent over-excitation and sustain the phasic neural activity of DRN_VIP_ neurons during risk assessment. Alternatively, it is likely that PKC-δ neurons in the CeA (lateral division) and ovBNST are heterogeneous, with divergent projections in each structure. Additional experiments will be necessary to distinguish between feedback loop versus divergent projection organization. Notably, we also demonstrated that DRN_VIP_ neurons occupy a central position, with the unique characteristic of individual neurons projecting to both the CeA (lateral division) and ovBNST (Supplementary Figure 4). In both regions, these neurons appear to exhibit coordinated activity and function as threat detectors in experimental paradigms that elicit defensive behaviors (Moscarello and Penzo, 2022; Williford et al., 2023). Collectively, these findings suggest that DRN_VIP_ neurons play a critical role in threat evaluation and the initiation of defensive responses.

### DRN_VIP_ neurons orchestrate active-phase sleep architecture, risk assessment and defensive behaviors

Our findings indicate that caspase-induced ablation of DRN_VIP_ neurons selectively alters NREM sleep architecture during the active-phase, prolonging individual sleep episodes while reducing their overall frequency. Notably, total NREM sleep time remained unchanged, ruling out excessive daytime sleepiness. This suggests that DRN_VIP_ neurons contribute to shorten the duration of sleep bouts, and conversely that their loss enhances sleep stability. Given that sleep stability is crucial for modulating cognitive and neural processes involved in threat evaluation and adaptive defensive behaviors (Loftus et al., 2022), DRN_VIP_ activity may serve as a regulatory mechanism balancing restorative sleep with wakefulness in dynamic environments. Previous work has demonstrated that DRN_DA_ neurons play a key role in arousal control, acting as a "gatekeeper" for wakefulness by promoting rapid awakening in response to salient stimuli (Cho et al., 2017). Our results suggest that DRN_VIP_ neurons could function as a neuronal alarm system, reacting to threats or unexpected stimuli—such as movement or noise from conspecifics—that are more likely to occur during the active phase. By maintaining NREM sleep episodes short, these neurons may ensure heightened vigilance when environmental disturbances are most frequent. Consequently, in their absence, increased sleep stability emerges specifically during the active phase without affecting overall homeostatic sleep need (Figure 5).

Our behavioral tests rely on an approach/avoidance conflict, balancing the drive to explore novel areas against aversion to potentially threatening environments. These tests inherently involve risk assessment and threat evaluation as the animal decides whether to enter a potentially unsafe area (Blanchard et al., 1993; Rodgers and Dalvi, 1997; Zhao et al., 2023). The EPM has been validated to quantify risk assessment and anxiety-like behaviors (Rodgers and Dalvi, 1997), with impaired risk assessment typically linked to increased time and more frequent entries into the open arms, reflecting heightened risk-taking behavior (Zhou et al., 2024). Our study found impaired risk assessment (Figure 8) behaviors in mice with genetic ablation of DRN_VIP_ neurons, consistent with the established role of DRN_DA_ neurons in detecting salient stimuli and assigning them positive or negative valence (Cho et al., 2021). Moreover, the BNST and CeA, primary targets of DRN_VIP_ neurons, are central to risk assessment, valence attribution, and defensive behaviors (Lebow and Chen, 2016). Furthermore, it has been shown in humans that changes in BNST and CeA connectivity during threat anticipation have been associated with anxiety and risk evaluation processes (Pedersen et al., 2019; Torrisi et al., 2018). Together, these findings underscore the pivotal role of DRN_VIP_ neurons and their targets, including PKC-δ neurons, in controlling risk assessment.

Overall, the genetic ablation of DRN_VIP_ neurons does not appear to affect the mice’s general anxiety state, except perhaps in the EPM, where lesioned mice spend less time in the open arms (Figure 6D). While this outcome may seem counterintuitive, our findings suggest that disrupting DRN_VIP_ neurons—a key component of the risk assessment circuit—impairs information gathering, reducing risky behavior in the EPM and discouraging exploration of riskier areas. The visual looming test is highly effective at triggering defensive behaviors while also offering valuable insights into exploration and risk assessment, making it a useful tool for studying impaired information gathering (Yilmaz and Meister, 2013). Here we show in the visual looming test, selective genetic ablation of DRN_VIP_ neurons leads to risk assessment and defensive behaviors impairment.

The looming test assesses the balance between exploration and safety, a critical component of risk assessment. Since risk assessment is a critical component in modulating escape behavior (Evans et al., 2019), the impaired escape behavior observed in mice lacking DRN_VIP_ neurons during the looming test could be influenced by an underlying maladaptive risk assessment behavior. Understanding which neural circuits transmit visual information to DRN_VIP_ neurons remains a key question. One potential answer lies in findings from mice, where retinal projections to the superior colliculus and the DRN are believed to play a key role in processing visual looming stimuli and mediating defensive behaviors (Huang et al., 2017; Shang et al., 2015; Wei et al., 2015). Our results reveal that neurons in the superior colliculus form synaptic connections with DRN_VIP_ neurons (Figure 2B), supporting the idea that the superior colliculus may serve as a relay for visual information to DRN_VIP_ neurons.

In conclusion, our study provides novel insights into the neural circuitry underlying sleep stability, risk assessment and defensive behaviors. Our findings underscore the importance of DRN_VIP_ neurons as central integrators of risk assessment circuits. The relevance of this circuitry extends beyond rodent models. Human studies link BNST and CeA connectivity to risk evaluation and anxiety processes, suggesting that similar neural mechanisms may underlie these behaviors across species. Importantly, our findings may provide valuable insights into autism spectrum disorder (ASD). Studies have shown that BNST synaptic functions are disrupted in an ASD mouse model (Contestabile et al., 2023), associated to abnormal risk-assessment behaviors (Oron et al., 2019). Additionally, a translational study reported that children with autism do not exhibit typical looming-evoked defensive responses (Hu et al., 2017). By elucidating the neural substrates of risk assessment, our study opens avenues for exploring targeted interventions in conditions like ASD, where these processes are disrupted.

## Supporting information

Table 1 - Key ressources

Table 2 - Statistics

## ACKNOWLEDGMENTS

This work has received funding from the European Union’s Horizon 2020 research and innovation program under grant agreement No 848002. This study received financial support from the French government in the framework of the University of Bordeaux’s IdEx "Investments for the Future" program / GPR BRAIN_2030. E.P. is a recipient of a Clément Fayat Foundation fellowship (France). This work was supported by recurrent funding from the University of Bordeaux and the CNRS. Support from the Swedish Research Council (Grant 2024-02966) to GF. We Thank Dr. Andreas Frick (Neurocentre Magendie, INSERM, U1215) for facilitating the transfer of the VIP-Cre mouse line from the Neurocentre Magendie animal facility to the PIV-EXPE at CBNA. We thank Aquineuro for helping us to develop the code for analyzing the defensive behaviors in the visual looming test. We thank Fabrice Cordelières for developing the ImageJ plugin (IJ-Plugin_Atlas-Utilities) used for mapping GPF fibers in cleared brains. The microscopy was done in the Bordeaux Imaging Center, part of CNRS-INSERM and Bordeaux University, member of the national infrastructure France BioImaging supported by the French National Research Agency (ANR-10-INBS-04).

## Author Contributions

Conceptualization: A.G.; C.G. G.F. and F.G. ; Investigation: A.G., F.G; E.P, S.D, S.D.; N.B.; E.L.; L.B.; D.D.C.M.; Resources: F.G., J.B., M.L. and E.B; Writing – Original Dra{: A.G. and F.G.; Visualization: A.G., T.D., E.P.; Supervision: F.G.; Funding Acquisition: F.G., J.B. and G.F.

## Resource Availability

Requests for further information and resources should be directed to and will be fulfilled by the lead contact, François Georges

## Materials availability

This study did not generate new unique reagents,

## Data and code availability

Code utilized to analyze the defensive behavior data are available from the lead contact upon request. Any additional information required to reanalyze the data reported in this paper is available from the lead contact upon request.

## STAR METHODS

### Animals

C57BL/6JRj (≥ 8 week-old; Elevage Janvier, France), Vip-IRES-Cre and Vip-IRES-Cre/COP4 mice were used. VIP-IRES-Cre and Vip-IRES-Cre/COP4 mice were maintained on a C57BL/6J genetic background. Animals were housed two to five per cage in standard temperature and humidity conditions (22-23°C, 40 % relative humidity, 12h light/dark illumination cycle) and had access to food and water *ad libitum.* Both female and male mice were used in this study. Female rhesus monkeys (Macaca mulatta: mean age = 5 ± 1years, mean weight = 5.3 ± 0.8 kg, n = 2) were euthanized with an overdose of pentobarbital (150 mg/kg). Brains were quickly removed after death, immediately frozen in dryice-cooled isopentane, and then stored at −80 °C. The time between euthanasia and freezing of the brains was 10 min in all cases.

### Double-probe fluorescent in situ hybridization (sdFISH)

sdFISH was carried out as described previously (Dumas et al., 2019).

Probes: sdFISH was performed using antisense riboprobes for the detection of Vip mRNA: NM_011702.3 sequence 402-1320, Cre: AB449974.1 sequence 245-1060 and Th: NM_012740.4 sequence 456-1453. Synthesis of digoxigenin and fluorescein-labeled RNA probes were made by a transcriptional reaction with incorporation of digoxigenin or fluorescein labelled nucleotides. Specificity of probes was verified using NCBI blast.

Procedure: C57BL/6J (N = 4) or Vip-Cre (N = 5) mice were sacrificed by cervical dislocation. Brains were frozen in -30 to -35°C cold 2-methylbutane (≥99%, Honeywell) and stored at -80°C until sectioned with cryostat (Leica). 16 µm thin cryosections were cut and directly mounted on glass slides in a serial way (9 slices of DRN distributed on 8 slides), air-dried and kept at - 80°C until used. Slices were fixed in 4% paraformaldehyde and acetylated in 0.25% acetic anhydride/100 mM triethanolamine (pH 8). Sections were hybridized for 18 h at 65°C in 100 µl of formamide-buffer containing 1 µg/ml digoxigenin-labeled riboprobe (DIG) and 1 µg/ml fluorescein-labeled riboprobe. Sections were washed at 65°C with SSC buffers of decreasing strength, and blocked with 20% fetal bovine serum and 1% blocking solution. Fluorescein epitopes were detected with horseradish peroxidase (HRP) conjugated anti-fluorescein antibody at 1:5000 and revealed using Cy2-tyramide at 1:250. HRP-activity was stopped by incubation of sections in 0.1 M glycine followed by a 3% H2O2 treatment. DIG epitopes were detected with HRP anti-DIG Fab fragments at 1:2000 and revealed using Cy3 tyramide at 1:100. Nuclear staining was performed with 4’ 6-diamidino-2-phenylindole (DAPI).

Data Analysis: All slides were scanned at 20x resolution using the Hamamatsu NanoZoomer S60 Digital slide scanner (Hamamatsu). Laser intensity and time of acquisition were set separately for each riboprobe. Images were analyzed using the NDP.view2 software (Hamamatsu Photonics). Regions of interest were identified according to the Paxinos mouse brain atlas. Positive cells refer to a staining in a cell body clearly above background and surrounding a DAPI-stained nucleus. Colocalization was determined by the presence of the signals for both probes in the soma of the same cell.

### Stereotaxic injections

Surgeries for anatomy, behavioral experiment and *in vivo* electrophysiology were performed as follow: mice were anesthetized with a mixture of air/isoflurane (maintained at 1-2% isoflurane, 1 min.L^-1^) and placed in a stereotaxic frame. After receiving subcutaneous doses of Meloxicam (20 mg/Kg) and Buprenorphine (0.1 mg/Kg) and a local injection of Lurocaine (7 mg/Kg) the skin was incised. Coordinates for targeting the following structures were: BNST, +0.15 mm from Bregma, -0.85 mm from sagittal vein and -3.85 mm from brain surface; CeA, - 1.65 mm from Bregma, -0.85 mm from sagittal vein and -3.85 mm from brain surface; DRN, -4.20 to -4.60 mm from bregma, -1.00 mm from sagittal vein and -2.80 for targeting VIP neurons or -3.00 for targeting the whole DRN dopaminergic neurons, angle: 20° in the coronal plane. Injected volumes were 3x300 nL for AAV-EF1-DIO-Caspase3 and and control injections in the DRN, 3x180 nL for AAV1-EF1A-DIO-synaptophysin-GFP, helper virus (AAV1-EF1a-DIO-TVA950-T2A-cvs11G-WPRE), modified helper virus (AAV1-EF1a-DIO-TVA950-T2A-WPRE) and tracing virus used for brain clearing (AAV5-EF1a-DIO-eYFP) in the DRN, 1x250 nL for rabies injections in DRN (PSADB19dG-mCherry) and 120 nL for rabies injections in ovBNST (PSADB19dG-GFP) and CeA (PSADB19dG-mCherry).

### Immunohistochemistry

Mice were first anesthetized with a mixture of isoflurane and air (4%, v/v isoflurane/air) and received s.c injection of Meloxicam followed 10 min later by i.p injection of lethal dose of Euthasol (300mg/Kg). After perfusion with 1X Phosphate-buffered saline (PBS), brains were extracted and post-fixed in 4% formaldehyde for at least 24 hours. 40 µm coronal brain slices were cut at the vibratome and transferred to 1X PBS wells. After 1X PBS washes an antigen retrieval step was performed for immunohistochemistries labelling VIP and PKC-δ. Afterwards, slices were incubated in a blocking solution for 90 min (5-10% normal goat serum in PBS-TritonX100 0.3% (S26-100ML, EMD Millipore Corp.). Directly after, slices were incubated in the primary antibody solution for 72h: for the histological analysis of the Casp3 behavioral group: guinea-pig anti-TH (1:2000, SYSY 213104) and rabbit anti-VIP (1:1000, ab 272726 Abcam); for control behavioral group (synaptophysin-GFP mice), chicken anti-GFP (1:1000, GFP-1020 Aves), guinea-pig anti-TH and rabbit anti-VIP. For modified rabies experiments, chicken anti-GFP, rat anti-RFP (1:1000, 5f8 Chromotek) and rabbit anti-VIP; for rabies experiments, mouse anti-PKC-δ (1:500, 610398 BD Biosciences), rat anti-RFP, rabbit anti-somatostatin (1:1000, NBP1-87022 Novus) for BNST and CeA or rabbit anti-2A peptide (1:1000, ABS31 Sigma), guinea-pig anti-TH for the DRN; for patch-clamp experiments, chicken anti-GFP, mouse anti-PKC-δ for BNST slices or chicken anti-GFP and rabbit anti-VIP for DRN slices. Slices were washed in 1X PBS and incubated in a secondary antibody solution (1:500 in 1X PBSTritonX-100) for all secondary antibodies for 2h containing antibodies matching the primary antibodies and desired fluorescent reporter listed here in the same order as above: goat anti-guinea-pig A568 (A11075 Life technologies), and goat anti-rabbit A647; goat anti-chicken A488, goat anti-guinea-pig A568 and goat anti-rabbit A647; goat anti-chicken A488, goat anti-rat A568 and goat anti-rabbit A647; goat anti-mouse A488, goat anti-rat A568, goat anti-rabbit A647; goat anti-rabbit A488, goat anti-guinea-pig A568; goat anti-chicken A488, streptavidin A568 (1:1000), goat anti-mouse A647; goat anti-chicken A488, streptavidin A568. Afterwards slices were washed in 1X PBS, incubated in Hoestch solution for 5 min (1:10000, 33258 Invitrogen), washed again and mounted on glass slides and coverslipped (Fluoromount-G, Sigma). Acquisitions were performed using either a Nanozoomer scanner (Hamamatsu), an epifluorescent microscope (Evident) or confocal microscope (Leica).

For non-human primate immunohistochemistry, double immunofluorescence staining was performed to visualize TH and VIP immunoreactivity simultaneously. Briefly, 50 µm free-floating sections were first washed in PBS and incubated in a blocking buffer containing 0.3% Triton X-100 and BSA (1/50 in 0.1 M PBS) to prevent nonspecific staining. The sections were then incubated overnight at room temperature with a mixture of primary antibodies diluted at 1 :1000 against TH and VIP (guinea pig anti-TH, Synaptic system; rabbit anti-VIP, Abcam).

After thorough rinsing in PBS, the sections were incubated for 1 hour at room temperature with a cocktail of corresponding secondary antibodies conjugated to either Alexa Fluor 488 or 568 fluorophores (Molecular Probes). Following PBS washes, the sections were mounted on gelatin-coated slides and incubated for 5 minutes in a 0.3% Sudan Black B/70% ethanol solution to reduce tissue autofluorescence. Counterstaining was performed by immersing the slides in Hoechst solution diluted in PBS for 5 minutes. Finally, the slides were coverslipped using a fluorescence mounting medium (Fluoromount-G, Sigma). Images were acquired using a Pannoramic Scanner II (3D Histech) at 20X magnification in extended mode, with five layers spaced 1.4 µm apart.

### Histological analysis

Behavioral experiment quantification: Slices containing DRN, ovBNST and CeA were used to quantify the amount of VIP and TH neurons (DRN) and fiber mean grey value (ovBNST and CeA) on images acquired with Leica confocal microscope SP5-MP at 20X magnification. Counting of VIP and TH neurons in the DRN was done manually using Fiji software. One slice of an intermediate level of the DRN was selected for each mouse. The mean grey value in the ovBNST and CeA was measured on 20 µm z-stack images using Fiji software.

Quantification of DRN_VIP_ neurons inputs: After immunolabelling for PKC-δ and amplifying rabies reporter with RFP antibody, images of coronal brain slices were acquired using either epifluorescent microscope (Evident) for quantifying whole brain inputs to DRNVIP neurons (xy mosaic images of whole slices at 10X magnification) or confocal microscope (SP5-MP, Leica) for quantifying colocalization of RFP and PKC-δ in the ovBNST and CeA (20X magnification). Whole brain quantification of DRN_VIP_ neurons was performed by using Fiji/QuPath softwares and an experimental plugin “Apposing Big brains and atlases” (ABBA) to apply the Allen Brain atlas classification to the brain slices images. RFP positive neurons were manually labelled on QuPath. Injection site (DRN) and closely surrounding regions were excluded from the quantification. Structures with a mean density of neurons below 5x10^-7^ were excluded from Figure 3B.

### Brain clearing

Brain clearing of the three brains has been done following Adipoclear protocol including several delipidation steps and GFP amplification (Chi et al., 2018). Whole brain acquisitions have been done using UltraMicroscope II from Biotech-Milteniy with a 1.26X magnification. Before whole brain quantification, fluorescence of injections sites has been measured and compared at 3 ventro-dorsal plans of the DRN per mouse (Supplementary Figure 3). Stack of acquisitions have been automatically aligned through Napari viewer following elastix registration to adapt the acquired brain to the Allen brain atlas. Counting of VIP+ cell bodies have been done using spots detection in Imaris software, with a diameter filter of 20um. Mean grey value quantification of each region of the Allen brain atlas has been done with a homemade plugin on Fiji. Whole brain quantification histograms have been obtained after min-max normalization of the dataset composed of brain regions with a signal intensity superior to 0.25% of total brain intensity (Figure 2).

### Ex vivo electrophysiological recording

Brain slice preparation: The brain was quickly removed from the skull and mounted on the stage of a vibratome (VT1200; Leica microsystems), immersed in a cutting solution at 4°C saturated with carbogen (N = 4 mice for ovBNST slices, N = 3 mice for CeA slices and N = 1 mouse for DRN slices). It was then sectioned into 300 μm thick slices in the coronal plane. Slices containing the ovBNST, CeA and DRN were then transferred to a holding chamber filled with standard carbogen-saturated ACSF and placed in a water bath at 32°C for 1 hour. The ACSF was composed of (in mM) NaCl (126), KCl (2.5), NaH2PO4·H2O (1.25), CaCl2·H2O (2), MgSO4·7H2O (2), NaHCO3 (26), D-glucose (10), glutathione (5), and sodium pyruvate (1). The pH of the ACSF was adjusted to 7.4 and its osmolarity between 310 and 315 mOsm. The slices were then maintained in this solution at room temperature until electrophysiological recordings.

Electrophysiology: The slices were then transferred one by one to the recording chamber of an upright microscope (AxioExaminer Z1; Zeiss) and continuously perfused with equilibrated ACSF heated to 32°C. In some experiments, AMPA/NMDA receptor antagonists, DNQX (50 µM) and D-APV (20 µM), were also used to demonstrate the glutamatergic nature of synaptic currents recorded in the ovBNST. Neurons were visualized using a 63X water immersion objective (APO-CHROMAT; Zeiss) with differential interference contrast (DIC) illumination or an epifluorescence system (HXP120; Kubler). Recordings of ovBNST neurons were made using patch electrodes made from borosilicate glass capillaries (GC150F10; Phymep) on a horizontal puller (Model P97; Sutter instruments) to obtain a pipette resistance of 5-8MΩ. In this project, the intracellular solution used is "KGluconate" which served for whole-cell recordings in current-clamp mode. It is composed of (in mM) KGluconate (135), NaCl (3.8), MgCl2 6H2O (1), HEPES (10), EGTA (0.1), Na2GTP (0.4), Mg1.5ATP (2). This solution also contains biocytin (2mg/ml) and has a pH of 7.25 and an osmolarity of 290-295 mOsm3. Electrophysiological signals were amplified and filtered with a 4KHz low-pass filter using a Multiclamp 700B amplifier (Molecular Devices) then digitized at 20 kHz (Digidata 1550, Molecular Devices) using Clampex 10.7 software (Molecular Devices). Whole-cell and voltage-clamp recordings were performed at a holding potential of -60 mV, and glutamatergic EPSCs evoked by optogenetic stimulation of VIP neurons in the DRN was measured. Their amplitude and kinetics were measured following a train of 10 light pulses at 10 hertz. After electrophysiological recordings, the slices were fixed overnight in a 4% paraformaldehyde (PFA) solution, then stored at 4°C in a 0.03% PBS-sodium azide solution until immunohistochemical processing.

### In vivo single-cell electrophysiological recordings

BNST and CeA neurons were recorded in Casp3 (N = 17 for ovBNST and N = 15 for CeA) and control mice (N = 15 for ovBNST and N = 5 for CeA). Mice are first anesthetized with a mixture of isoflurane, air and oxygen (4% isoflurane for anesthesia induction and maintained at 1-1.5% isoflurane 1 min.L^-1^ 50% air/oxygen). Mice were placed on a stereotaxic frame and a subcutaneous injection of a local analgesic (lidocaine, 7 mg/Kg) was done at the skin incision location. A craniotomy was made above the BNST and CeA on the right hemisphere. Recording glass micropipette (GC150F-10, Harvard Apparatus) was filled with 2% pontamine sky blue solution in 0.5 M sodium acetate and lowered into either BNST or CeA. Single-cell extra-cellular recordings were performed in these two structures for Casp3 mice and controls. Single-neuron spikes were amplified with an Axoclamp2B (300 Hz / 0.5 kHz), filtered (AM system) and collected online (CED 1401, Spike2, Cambridge Electronic Design). Validation of the correct location of the recordings was done a posteriori from the pontamine sky blue spot.

### Electrode implant for sleep recording

After AAV-EF1-DIO-Caspase3 and control injections in DRN, Vip-Cre mice (N=8 per group) were implanted with micro-screws over the left parietal cortex (AP −2.0 mm, ML 1.8 mm from bregma) for cortical activity recordings and the occipital bone as a reference. Stainless-steel electrodes with Teflon insulation (Model 791400, A-M Systems Inc., Carlsborg, WA, USA) were implanted in the neck muscles to record electromyographic (EMG) signals. The electrodes and micro-screws were then soldered onto a Straight Male PCB Header-4 connector, which was secured to the skull with dental cement and acrylic.

### Polysomnographic and Behavioral Data Acquisition

EEG and EMG signals were amplified at a 5000× gain and processed with a bandpass filter— applying a 1 Hz high-pass for all channels and setting low-pass filters at 3 kHz for EEG and 500 Hz for EMG—using signal conditioners (ERS100C and EMG 100 C amplifiers, Biopac MP160). These signals were digitized at a 6250 Hz sampling rate via AcqKnowledge software (v4.1, Biopac Systems) and stored for subsequent offline analysis. Each mouse underwent two consecutive recording sessions over a three-day period. On the first day, animals were acclimated to the recording environment and the apparatus from 10:00 to 16:00 h (the light-on, inactive phase). Recording started the following day at 18:00 h and continued until 19:00 h the next day. The complete 24-hour recording cycle consisted of 12 h of lights-off (from 19:00 to 7:00 h, corresponding to the active period) followed by 12 h of lights-on (from 7:00 to 19:00 h, representing the inactive period), with the initial hour (from 18:00 to 19:00 h) discarded to minimize stress-induced artifacts. For each session, four freely moving mice from the same home cage were placed in individual cages outfitted with bedding, nesting material, food, and water. Behavioral activity during polysomnography was captured using 1080p Logitech Brio webcams, and the EEG, EMG, and behavioral data were merged into a single video file at 10 frames per second using OBS Studio (v27.0.1).

### State sorting

The 24-hour polysomnography and behavioral recording period was manually divided into 10-second segments, each classified as AWAKE, NREM or REM sleep. AWAKE was identified by low-amplitude, desynchronized EEG signals paired with continuous EMG activity, whether during movement or rest. NREM sleep was characterized by an increase in EEG amplitude dominated by slow waves, low EMG tone, and a lack of movement. In contrast, REM sleep was marked by immobility, consistent theta-dominated EEG patterns, and virtually no EMG activity. The placement of a cortical electrode over the dorsal hippocampus enabled optimal detection of theta oscillations and distinction between REM and NREM stages. This integrated approach allowed for accurate sleep state classification and a detailed analysis of sleep architecture.

### Analysis of Sleep Architecture and EEG Microstructure

A custom MATLAB script was used to analyze sleep architecture and cortical activity. The duration of AWAKE, NREM, and REM states was calculated over the full 24-hour cycle by segmenting data into 1-hour bins and then averaging these values over 12-hour periods based on the light conditions (19:00–07:00 for lights off and 07:00–19:00 for lights on). NREM sleep fragmentation was evaluated in 1-hour bins by counting the number and measuring the duration of sleep bouts, with the results averaged over 12-hour periods.

Cortical EEG signals were processed using standard MATLAB functions. Initially, a digital low-pass infinite impulse response (IIR) filter (implemented with MATLAB’s designfilt function set to a 625 Hz cutoff) was applied to limit the frequency range, and the signal was then down-sampled to 1250 Hz. The signal was divided into 10-second segments, and for each epoch, the power spectral density was estimated using the pwelch function (with no overlap and 1 Hz steps). The power for each frequency bin was normalized by dividing by the total power of that epoch, and these normalized values were categorized into specific frequency bands: delta (1–4 Hz), theta (6–9 Hz), sigma (10–15 Hz), beta (16–30 Hz), and gamma (40–80 Hz). Each 10-second EEG segment was subsequently classified as AWAKE, NREM or REM sleep based on manual scoring.

Spindle analysis was performed with a custom MATLAB script (Medeiros et al., 2023). In brief, contiguous NREM EEG segments were filtered using a bandpass IIR filter (via designfilt) to isolate the spindle sigma frequency range. The signal envelope was then computed using a 700 ms moving root-mean-square (RMS) window, and the RMS value was cubed. Spindles were identified as events exceeding twice the cubed RMS average (calculated over the entire NREM period) with durations between 0.5 and 5 seconds. Finally, spindle density (number of events per minutes) and duration (in seconds) were quantified across the full 24-hour cycle during NREM sleep.

### Anxiety-related behavioral tests

Casp3 and control mice performed an open field test (OFT), a light/dark box test (L/D box) and an elevated plus maze test (EPM) described below. Prior to the first test, mice were handled progressively for three days. Before each test mice were habituated to the experimental room for 30 min. Video acquisition is done using Polyvision software (Imetronics) and video analysis by using Ethovision XT17 (Noldus). Number of mice for L/D box and EPM tests was N = 19 controls and N = 19 Casp3 mice and number of mice for the OFT was N = 17 controls and N = 20 Casp3 mice.

*Open field test*: the test consists of a square arena (40x40x40) where the mouse can freely explore for 10 min. The arena is artificially divided into zones: center, borders and corners. Traveled distance, velocity and time in the different zones were measured. The ROUT method was used on the “Time spent in center” variable to exclude potential outliers.

Light/Dark box: The arena is composed of two equally sized compartments (20x20x20 for each compartment), a light compartment strongly lit up (900-1000 lux) and a dark compartment. Both compartments are connected through a little aperture. The mouse is placed in the light compartment and can freely explore the arena for 10 min. Time spent in compartments, latency to light compartment, entries to the light compartment and nose pokes to the light compartment were measured. ROUT method was used on the “time spent in the light compartment” variable to exclude potential outliers.

*Elevated plus maze*: the apparatus consists in an elevated plus shape platform with two closed arms (25 cm walls) and two open arms (30 cm long, 5 cm wide). The mouse is placed in the center facing an open arm and can freely explore the arena for 10 min. Measured variables were time spent in open arms, closed arms and center, protected and unprotected headips and stretch attend postures (SAP) as well as rearing. Protected zones correspond to the closed arms and center area while unprotected zones correspond to the open arms. ROUT method was used on the “time spent in open arms” variable to exclude potential outliers.

*Visual looming test:* the visual looming test (VLT) setup consists in a transparent rectangular arena (36x19.7x20cm) divided in three zones: a shelter zone (12.7x19.7 cm) which has a roof over, an intermediate zone (13.3x19.7 cm) and a trigger zone (9.9x19.7 cm). A screen was placed about 30 cm above the ground of the cage with a light grey background. Mice (N = 19 controls and N = 19 Casp3 mice) had three days of habituation to the apparatus, 5 min every time (one day of collective habitation two to three mice together and two days of individual habituation). On day 1, the mouse was placed in the trigger zone and can freely explore the arena during 5 min. After 5 min of habituation (corresponding to H1 for Day1, H2 for Day2 and H7 for Day7), a visual cue, which corresponds to a trial, was manually triggered on the screen every time the mouse entered the trigger zone. A refractory period of 30 s was set after each visual cue. The test was considered completed either when the mouse performed 10 trials or after 60 min. The same protocol was done at day 2 and day 7. The visual cue consists in an expending black round which appears on the light grey background of the screen, first slowly from 0.5 to 3 cm then two fast times from 3 cm to 21 cm. The whole duration of the visual cue lasted for 6 s. The test was filmed using a GoPro camera and analyzed with DeepLabCut and python scripts (Aquineuro) for the following parameters: escape probabilities, time spent in zones, latency to escape, distance traveled, velocity and immobility were measured and manually for the rearing. Trials had to fulfill three criteria to be considered as escaping trials: a speed above 35 cm/s, a head orientation towards the shelter and the mouse position in the shelter in at least 7 s after the onset of the visual cue. Escape probability corresponds to the number of escapes divided by the number of trials. A mouse was considered immobile when velocity was below 1 cm/s for at least 1 s. Additionally, the vigor of the “escape” was measured by lowering the velocity threshold of the “escape” to 20 cm/s in order to measure the vigor of the return to the shelter. For rearing quantification, the before cue period includes 30 s before trial 3, 4, 5, 6 and 7 while during cue includes 30 s after trials 1, 2, 3, 4 and 5. ROUT method was used on the “rearing” variable to exclude potential outliers.

### Sleep, risk Assessment, defensive behavior, anxiety and locomotion z-score calculation

To assess behavioral dimensions such as sleep, risk assessment, defensive behavior, anxiety, and locomotion, we employed the integrated behavioral z-scoring method as described by (Guilloux et al., 2011). This approach standardizes individual behavioral measures by calculating z-scores, which represent the number of standard deviations a data point deviates from the mean of a control group. Specifically, for each behavioral parameter, the raw data were transformed into z-scores using the formula: z = (X - μ) / σ, where X is the individual score, μ is the mean of the control group, and σ is the standard deviation of the control group.

For the calculation of the NREM Sleep score during the active and inactive phases, the following raw data were used: time spent in NREM sleep, spindle duration during NREM, spindle density during NREM, and bout length during NREM sleep. The risk assessment score was derived from the percentage of protected stretch-attend postures (SAP) in the elevated plus maze (EPM), the number of nose pokes in the light-dark box (LDB), the number of rearings in the visual looming test (VLT), and the percentage of protected head dips in the EPM.

Defensive behavior was assessed using raw data from the VLT, including escape probability, maximum escape speed, immobility duration, and latency to escape. The anxiety score was calculated using the percentage of time spent in the open arms of the EPM, latency to enter the light compartment in the LDB, number of entries into the light compartment in the LDB, total time spent in the light compartment in the LDB, percentage of time spent in the center in the open field test (OFT), and time spent in the trigger zone during habituation on day 2 of the VLT. Lastly, the locomotion score was derived from total distance moved in the VLT during habituation on day 1, distance moved in the LDB, number of entries into the closed arms of the EPM, total distance traveled in the EPM, and distance moved in the OFT.

### Quantification and statistical analysis

We performed the statistical analysis using GraphPad Prism 5 (GraphPad, LaJolla, CA). Sample size (n) generally represents the number of experimental replicates, as indicated in the figure legends. In all behavioral experiments, (n) refers to the number of mice used, while in slice recording experiments, (n) denotes the number of recorded cells. Data are presented as mean ± SEM in all figures. For two group comparisons, statistical significance was determined by two-tailed paired or unpaired Student’s t-tests or non-parametric analogs, when assumptions for parametric testing were not satisfied. Normality was tested using D’Agostino and Pearson omnibus normality test. For multiple group comparisons, one-way analysis of variance (ANOVA) tests were used for normally distributed data, followed by post hoc analyses. For data that were not normally distributed, non-parametric tests for the appropriate group types were used instead, such As Mann-Whitney. Exact p-values and the corresponding statistical methods are provided in the figure captions and legends. A detailed description of the statistical tests can be found in Table 2.

### Animals and ethics approval

All animal procedures were conducted European directive 2010-63-EU and with approval from the Bordeaux University Animal Care and Use Committee (license authorization 21134). All efforts were made to minimize animal suffering and reduce the number of animals used. Non-human primate experiments were performed on tissue from a previously published brain bank (Fernagut et al., 2010; Fridjonsdottir et al., 2021; Porras et al., 2012). Experiments were carried out in accordance with the European Communities Council Directive of November 24, 1986 (86/609/EEC) regarding care of laboratory animals in an AAALAC-accredited facility following acceptance of the study design by the Institute of Lab Animal Science (Chinese Academy of Science, Beijing, China) IACUC. All procedures adhered to the 3Rs principles (Replacement, Reduction, and Refinement) to ensure ethical and humane research practices.

## KEY RESOURCE TABLE

**Table 1. Key resources table.**

**Table 2. Summary of statistical analyses.**

## SUPPLEMENTAL INFORMATION TITLES AND LEGENDS

**Supplementary Figure 1. *In situ* hybridization of Vip and Th mRNA in the DRN in wild-type (C57BL/6J) and Vip-Cre mice. (A)** Fluorescent images of Th and Vip mRNAs in the DRN in C57BL/6J mice associated to a pie chart showing co-localization quantification. (B) Fluorescent images of Th and Vip mRNAs in the DRN in Vip-Cre mice on the top associated to a pie chart showing co-localization quantification. Below, fluorescent images of Cre recombinase and Vip mRNAs in the DRN in Vip-Cre mice associated to a pie chart showing co-localization quantification. Th: tyrosine hydroxylase; Vip: vasoactive intestinal peptide; Cre: Cre recombinase; aq: aqueduct of Sylvius.

**Supplementary figure 2. Characterization of DRN_VIP_ neurons and terminals in the non-human primate (NHP). (A)** Schematics of a NHP and colorimetric immunostaining of VIP in the DRN (in brown) with a Nissl staining (purple). **(B)** Double immunofluorescence of VIP and TH in the DRN in the NHP with closeup images showing colocalization of VIP and TH. **(C)** Colorimetric immunostaining of VIP fibers in the ovBNST and CeA in the NHP. **(D)** Close-up images of VIP fibers in the ovBNST (on the top) and CeA (on the bottom) from C. **(E)** On the left, overview brightfield image of one hemisphere at the level of the ovBNST with a Black Sudan staining. On the right, same image with TH fluorescent immunostaining associated to close-up fluorescent images of the ovBNST with TH and VIP immunostaining. **(F)** On the left, overview brightfield image of one hemisphere at the level of the CeA with a Black Sudan staining. On the right, same image with TH fluorescent immunostaining associated to close-up fluorescent images of the CeA with TH and VIP immunostaining. Abbreviations: VIP: vasoactive intestinal peptide; TH: tyrosine hydroxylase; aq: aqueduct of Sylvius; DRN: dorsal raphe nucleus; ovBNST: oval nucleus of the bed nucleus of the stria terminalis; Cd: caudate putamen; Pu: putamen; GPe: external globus pallidus; GPi; internal globus pallidus; CeA: central nucleus of the amygdala; Thal: thalamus.

**Supplementary figure 3. Brain clearing and quantification approach of DRNVIP projections. (A)** Whole brain acquisition of autofluorescence (in grey) and DRN VIP cell bodies (in red). **(B)** Comparison of intensity level in DRN injection sites (Brown-Forsythe test, p-value = 0.9579). **(C, E)** Mean signal intensity of the three brains at DRN/ST terminalis **(C)** and CeA/BNST level **(E)**. **(D, F)** Mean heatmaps of labeling with atlas annotations. Scale bars: 1000 µm.

**Supplementary figure 4. Mapping of DRN_VIP_ collaterals to ovBNST and CeA. (A)** Schematic representation of the viral strategy used to demonstrate DRN_VIP_ collaterals to ovBNST and CeA. For this experiment a modified helper virus lacking the glycoprotein (AAV1-EF1a-DIO-TVA950-T2A-WPRE) was used associated with pseudorabies viruses conjugated to two different fluorescent reporters. **(B)** Confocal images of DRN neurons expressing either one or two of the pseudorabies reporters. On the top are neurons located near the aqueduct of Sylvius and below are DRN neurons located more ventrally.

**Supplementary figure 5. Validation of the rabies strategy used for tracing inputs to DRN_VIP_ neurons. (A)** Schematic representation of the helper virus (AAV1-DIO-TVA950-T2A-csv11G) and pseudorabies virus (pSADB19dG-mCherry) injections in the DRN in Vip-Cre mice. **(B)** Confocal images of starter cells with colocalization of T2A, RFP and TH in helper cells.

**Supplementary Figure 6. Effect of DRN_VIP_ genetic ablation on AWAKE and REM states in Vip-Cre mice. (A)** Graphics showing AWAKE percent time during the active (19- to 7 h) and the inactive (7- to 19 h) phases in Ctrl and Casp3 groups. **(B, C)** Bar graphs showing the AWAKE percent time during the active (19- to 7 h), (B, Unpaired T-test p-value=0.4488) and the inactive (7- to 19 h), (C, Unpaired T-test p-value=0.7368) phases in Ctrl and Casp3 groups. **(D, E)** Bar graphs showing the number (D, Unpaired T-test p-value=0.0232) and the length (E, Unpaired T-test p-value=0.1809) of AWAKE bouts during the active phase (19- to 7 h) in Ctrl and Casp3 groups. **(F, G)** Bar graphs showing the number (F, Mann-Whitney p-value=0.4889) and the length (G, Unpaired T-test p-value=0.1949) of AWAKE bouts during the inactive phase (7- to 19 h) in Ctrl and Casp3 groups. **(H)** Graphics showing REM percent time during the active (19- to 7 h) and the inactive (7- to 19 h) phases in Ctrl and Casp3 groups. **(I, J)** Bar graphs showing the REM sleep percent time during the active (19- to 7 h), (I, Unpaired T-test p-value=0.6160) and the inactive (7- to 19 h), (J, Unpaired T-test p-value=0.6850) phases in Ctrl and Casp3 groups. **(K, L)** Bar graphs showing the number (K, Unpaired T-test p-value=0.3492) and the length (L, Unpaired T-test p-value=0.0521) of REM sleep bouts during the active phase (19- to 7 h) in Ctrl and Casp3 groups. **(M, N)** Bar graphs showing the number (M, Unpaired T-test p-value=0.2228) and the length (N, Unpaired T-test p-value=0.5133) of REM sleep bouts during the inactive phase (7- to 19 h) in Ctrl and Casp3 groups. Light circles in graphs correspond to female mice while dark circles correspond to male mice. *p-value < 0.5.

**Supplementary figure 7.** Impact of DRN_VIP_ neurons genetic ablation on i*n vivo* electrophysiological properties of BNST and CeA neurons. (A) Schematic representation of AAV5-EF1a-Flex-Casp3 or control virus injections in the DRN in Vip-Cre mice. **(B)** Graphs showing the firing rate of BNST and CeA neurons in Casp3 and control mice (BNST, Mann-Whitney p-value=0.0045); CeA, Mann-Whitney p-value=0.0227). **(C)** Example of brightfield and fluorescent images of the last recorded coordinates in the BNST and CeA shown by the injection of pontamine sky blue (in red).

