## Supplementary material for "A subset of dorsal raphe dopamine neurons is critical for survival-oriented vigilance": Table 1 - Key ressources

Supplementary table 1

| Reagent or resource | Source | Identifier |
|---|---|---|
| Antibodies | | |
| Rabbit anti-VIP | Abcam | Cat. #ab272726 |
| Mouse anti-PKC-δ | BD Biosciences | Cat. #610398; RRID: AB_397781 |
| Guinea-pig anti-TH | Synaptic system | Cat. #213104; RRID:AB_2619897 |
| Mouse anti-TH | Milipore | Cat. #MAB318; RRID: AB_2201528 |
| Chicken anti-GFP | Aves Lab | Cat. #GFP-1020; RRID: AB_10000240 |
| Rat anti-RFP | Chromotek | Cat. #5f8; RRID: AB_2336064 |
| Rabbit anti-2A peptide | Sigma-Aldrich | Cat. #ABS31; RRID: AB_11214282 |
| Goat anti-guinea-pig A568 | Life technologies | Cat. #A11075; RRID: AB_141954 |
| Goat anti-mouse A488 | Invitrogene | Cat. #A11001; RRID: AB_2534069 |
| Goat anti-chicken A488 | Invitrogene | Cat. #A11039; RRID: AB_2534096 |
| Goat anti-rabbit A488 | Invitrogene | Cat. #A11008; RRID: AB_143165 |
| Goat anti-rat A568 | Invitrogene | Cat. #A11077; RRID: AB_2534121 |
| Goat anti-guinea-pig A568 | Life technologies | Cat. #A11075; RRID: AB_141954 |
| Goat anti-mouse A647 | Life technologies | Cat. #A21236; RRID: AB_2535805 |
| Goat anti-rabbit A647 | Life technologies | Cat. #A21245; RRID: AB_2535813 |
| **Viral vectors** | | |
| AAV1-EF1a-FLEX-synaptophysine | IMN VectorCore | |
| AAV5-EF1a-DIO-EYFP | Addgene | Cat. #v147551 |
| pSADB19dG-mCherry | Viral Core Facility, Charité - Universitätsmedizin Berlin | Cat. #RV03 |
| pSADB19dG-GFP | Viral Core Facility, Charité - Universitätsmedizin Berlin | Cat. #RV03 |
| AAV1-EF1a-DIO-TVA950-T2A-WPRE | IMN VectorCore | Cat#PV351 |
| AAV1-EF1a-DIO-TVA950-T2A-cvs11G-WPRE | IMN VectorCore | |
| AAV5-pAAV-flex-taCasp3-TEVp | Addgene | v143243// 45580-AAV5 |
| **Chemicals** | | |
| Streptavidin A568 | Life technologies | Cat#S11226, RRID: AB_2315774 |
| Normal goat serum | Sigma-Aldrich | S26-100ML |
| Hoechst | Invitrogene | RRID: AB_2651133 |
| DAPI | Invitrogene | Cat. D1306, RRID:AB_2629482 |
| DNQX disodium salt | Tocris | Cat. #2312 |
| D-AP5 | Tocris | Cat. #0106 |
| Pontamine sky blue | Sigma-Aldrich | Cat. #C8679 |
| Euthasol | centravet | |
| Meloxicam | centravet | |
| Lurocaine | centravet | |
| Buprenorphine | virbac | |
| Isoflurane | virbac | |
| **Experimental models: Organisms/strains** | | |
| C57BL/6JRj | Janvier Labs | SC-C57J-F and M |
| Vip-IRES-Cre | On site production (PIV-EXPE) | RRID:IMSR_JAX:010908 ; Strain #:010908 |

| | | |
|---|---|---|
| Vip-IRES-Cre/COP4 | On site production (PIV-EXPE) | COP4: RRID:IMSR_JAX:024109 ; Strain #:024109 |
| Software and algorithms | | |
| Spike2 | Cambriddge Electronic Design Limited | RRID: SCR_000903 |
| ImageJ | | RRID:SCR_003070 |
| Fiji | | RRID: SCR_002285 |
| Qupath v5.0 | Bankhead, P. et al. 2017 | RRID: SCR_018257 |
| Abba | BioImaging & Optics Platform, EPFL | Atlas V3p1; RRID: SCR_023857 |
| Imaris | Oxford Instruments | RRID: SCR_007370 |
| Ethovision XT v17.0 | Noldus | RRID: SCR_004074 |
| Deeplabcut | Deeplabcut | RRID: SCR_021391 |
| Graphpad Prism | Graphpad Prism | RRID: SCR_002798 |
| NDP.view2 | Hamamatsu | RRID: SCR_025177 |
| Inkscape | Inkscape | RRID: SCR_014479 |
| Imaging systems | | |
| Epifluorescent microscope | Olympus BX63 | |
| Confocal microscope | Leica TCS SP5 | |
| Digital Slide Scanner | Hamamatsu Nanozoomer 2.0HT | |
| UltraMicroscope II | Biotech-Milteniy | |
