## Supplementary material for "A subset of dorsal raphe dopamine neurons is critical for survival-oriented vigilance": Table 2 - Statistics

| Figure | Test type | Compared groups | Measured variable (Graph title) | Statistic test | Source of variation | P-value |
|---|---|---|---|---|---|---|
| Figure 4C | Histological quantification | Ctrl vs. Casp3 | VIP+ neurons DRN | Mann-Whitney | | **<0.0001** |
| Figure 4C | Histological quantification | Ctrl vs. Casp3 | TH+ neurons DRN | Unpaired T-test | | 0.0753 |
| Figure 4D | Histological quantification | Ctrl vs. Casp3 | VIP+ fibers ovBNST | Mann-Whitney | | **<0.0001** |
| Figure 4D | Histological quantification | Ctrl vs. Casp3 | TH+ fibres ovBNST | Mann-Whitney | | **<0.0001** |
| Figure 4E | Histological quantification | Ctrl vs. Casp3 | VIP+ fibers CeA | Mann-Whitney | | **<0.0001** |
| Figure 4E | Histological quantification | Ctrl vs. Casp3 | TH+ fibres CeA | Mann-Whitney | | **<0.0001** |
| Figure 5E | Sleep | Ctrl vs. Casp3 | NREM time active (%) | Unpaired T-test | | 0.4375 |
| Figure 5F | Sleep | Ctrl vs. Casp3 | NREM time inactive (%) | Unpaired T-test | | 0.7605 |
| Figure 5H | Sleep | Ctrl vs. Casp3 | NREM bouts active (n) | Unpaired T-test | | **0.0207** |
| Figure 5I | Sleep | Ctrl vs. Casp3 | NREM bouts duration active | Unpaired T-test | | **0.0038** |
| Figure 5J | Sleep | Ctrl vs. Casp3 | NREM bouts inactive (n) | Unpaired T-test | | 0.3486 |
| Figure 5K | Sleep | Ctrl vs. Casp3 | NREM bouts duration inactive | Unpaired T-test | | 0.3050 |
| Figure 5L | Sleep | Ctrl vs. Casp3 | Spindles density active | Unpaired T-test | | 0.2357 |
| Figure 5M | Sleep | Ctrl vs. Casp3 | Spindles duration active | Unpaired T-test | | **0.0118** |
| Figure 5N | Sleep | Ctrl vs. Casp3 | Spindles density inactive | Unpaired T-test | | 0.5951 |
| Figure 5O | Sleep | Ctrl vs. Casp3 | Spindles duration inactive | Unpaired T-test | | 0.6303 |
| Figure 5C | OFT | Ctrl vs. Casp3 | Time spent in center (%) | Unpaired T-test | | 0.3591 |
| Figure 6C | OFT | Ctrl vs. Casp3 | Distance moved (cm) | Unpaired T-test | | 0.4645 |
| Figure 6D | EPM | Ctrl vs. Casp3 | OA/(OA+CA) *100 (%) | Unpaired T-test | | **0.0267** |
| Figure 6D | EPM | Ctrl vs. Casp3 | Protected Headips (%) | Unpaired T-test | | **0.0396** |
| Figure 6D | EPM | Ctrl vs. Casp3 | Protected SAP (%) | Mann-Whitney | | 0.2841 |
| Figure 6D | EPM | Ctrl vs. Casp3 | Time spent in center (%) | Unpaired T-test | | 0.8845 |
| Figure 6D | EPM | Ctrl vs. Casp3 | Nb of entries in CA | Unpaired T-test | | 0.7502 |
| Figure 6D | EPM | Ctrl vs. Casp3 | Distance moved (cm) | Unpaired T-test | | 0.329 |
| Figure 6E | L/D box | Ctrl vs. Casp3 | Time in light (%) | Unpaired T-test | | 0.9668 |
| Figure 6E | L/D box | Ctrl vs. Casp3 | Nb of entries in light | Unpaired T-test | | 0.7103 |
| Figure 6E | L/D box | Ctrl vs. Casp3 | Latency to light (s) | Unpaired T-test | | 0.2553 |
| Figure 6E | L/D box | Ctrl vs. Casp3 | Distance moved in light (cm) | Unpaired T-test | | 0.9202 |
| Figure 6E | L/D box | Ctrl vs. Casp3 | Nb of nose pokes | Unpaired T-test | | 0.8581 |
| Figure 7D | VLT | Ctrl vs. Casp3 (Group) and Days | Escape probability along days | Mixed-effects analysis | $F_{Group}$ (1, 32) = 10.20 | **0.0031** |
| Figure 7D | VLT | Ctrl vs. Casp3 | Escape probability Day1 | Mann-Whitney | | **0.0013** |
| Figure 7D | VLT | Ctrl vs. Casp3 | Escape probability Day2 | Mann-Whitney | | **0.0423** |
| Figure 7D | VLT | Ctrl vs. Casp3 | Escape probability Day7 | Mann-Whitney | | **0.0034** |
| Figure 7E | VLT | Ctrl vs. Casp3 | Donuts percentage escaping/no escaping Day1 | Fisher's exact test (Contingency) | | **<0.0001** |

| | | | | | | |
|---|---|---|---|---|---|---|
| Figure 7F | VLT | Ctrl vs. Casp3 | Max velocity during cue Day1 (cm/s) | Mann-Whitney | | **0.0056** |
| Figure 7G | VLT | Ctrl vs. Casp3 (Group) and Days | Latency 1st escape along days | Mixed-effects analysis | $F_{Days}$ (1.928, 53,03) = 7.372 | **0.0017** |
| Figure 7H | VLT | Ctrl vs. Casp3 (Group) and Days | Time in shelter along days (%) | 2Way RM ANOVA | $F_{Group}$ (1, 32) = 5.343, $F_{Days}$ (1.892, 60,53) = 3.747 | **Group: 0.0274, Days: 0.0314** |
| Figure 7H | VLT | Ctrl vs. Casp3 (Group) and Days | Time in trigger zone along days (%) | 2Way RM ANOVA | $F_{Group}$ (1, 32) = 4.616 | **0.0393** |
| Figure 7I | VLT | Ctrl vs. Casp3 | Time in trigger zone H2 (%) | Mann-Whitney | | 0.4174 |
| Figure 7J | VLT | Ctrl vs. Casp3 | Nb of rearing H1 | Mann-Whitney | | 0.4375 |
| Figure 7J | VLT | Ctrl vs. Casp3 | Rearing (% from H1) Before cue | Unpaired T-test; One sample t test | | **0.0062;** 0.3974 (Ctrl)**, 0,0003 (Casp3)** |
| Figure 7J | VLT | Ctrl vs. Casp3 | Rearing (% from H1) After cue | Mann-Whitney; One sample t test | | **0.0046; 0.0013 (Ctrl),** 0.6910 (Casp3) |
| Figure 7K | VLT | Ctrl vs. Casp3 | Cumulated immobility Day1 (s) | Unpaired T-test | | **0.0135** |
| Figure 7L | VLT | Ctrl vs. Casp3 | Distance moved H1 (cm) | Mann-Whitney | | 0.6212 |
| Figure 8A | Z-Score NREM (active) | Ctrl vs. Casp3 | Z-Score NREM active | Unpaired T-test | | **0.0012** |
| Figure 8B | Z-Score NREM (inactive) | Ctrl vs. Casp3 | Z-Score NREM inactive | Unpaired T-test | | 0.7542 |
| Figure 8C | Z-Score Risk assessment | Ctrl vs. Casp3 | Z-Score Risk assessment | Unpaired T-test | | **0.0075** |
| Figure 8D | Z-Score Defensive Behavior | Ctrl vs. Casp3 | Z-Score Defensive Behavior | Mann-Whitney | | **0.0009** |
| Figure 8E | Z-Score Anxiety | Ctrl vs. Casp3 | Z-Score Anxiety | Mann-Whitney | | 0.4654 |
| Figure 8F | Z-Score Locomotion | Ctrl vs. Casp3 | Z-Score Locomotion | Mann-Whitney | | 0.5334 |
| Supplementary Figure 3B | Brain clearing | AAV5-EF1-DIO-eYFP injected brains (DRN) | Mean grey value in DRN | Brown-Forsythe ANOVA test | | 0.9579 |
| Supplementary Figure 6B | Sleep | Ctrl vs. Casp3 | AWAKE time active (%) | Unpaired T-test | | 0.4488 |
| Supplementary Figure 6C | Sleep | Ctrl vs. Casp3 | AWAKE time inactive (%) | Unpaired T-test | | 0.7368 |
| Supplementary Figure 6D | Sleep | Ctrl vs. Casp3 | AWAKE bouts active (n) | Unpaired T-test | | **0.0232** |
| Supplementary Figure 6E | Sleep | Ctrl vs. Casp3 | AWAKE bouts duration active | Unpaired T-test | | 0.1809 |
| Supplementary Figure 6F | Sleep | Ctrl vs. Casp3 | AWAKE bouts inactive (n) | Unpaired T-test | | 0.4889 |

| | | | | | | |
|---|---|---|---|---|---|---|
| Supplementary Figure 6G | Sleep | Ctrl vs. Casp3 | AWAKE bouts duration inactive | Mann-Whitney | | 0.1949 |
| Supplementary Figure 6B | Sleep | Ctrl vs. Casp3 | REM time active (%) | Unpaired T-test | | 0.6160 |
| Supplementary Figure 6C | Sleep | Ctrl vs. Casp3 | REM time inactive (%) | Unpaired T-test | | 0.6850 |
| Supplementary Figure 6D | Sleep | Ctrl vs. Casp3 | REM bouts active (n) | Unpaired T-test | | 0.3492 |
| Supplementary Figure 6E | Sleep | Ctrl vs. Casp3 | REM bouts duration active | Unpaired T-test | | 0.0521 |
| Supplementary Figure 6F | Sleep | Ctrl vs. Casp3 | REM bouts inactive (n) | Unpaired T-test | | 0.2228 |
| Supplementary Figure 6G | Sleep | Ctrl vs. Casp3 | REM bouts duration inactive | Unpaired T-test | | 0.5133 |
| Supplementary Figure 7B | *In vivo* electrophysiology | Ctrl vs. Casp3 | Firing rate (BNST) | Mann-Whitney | | **0.0045** |
| Supplementary Figure 7B | *In vivo* electrophysiology | Ctrl vs. Casp3 | Firing rate (CeA) | Mann-Whitney | | **0.0227** |
