## Supplementary figures and images for "A subset of dorsal raphe dopamine neurons is critical for survival-oriented vigilance"

### Supplementary Figures

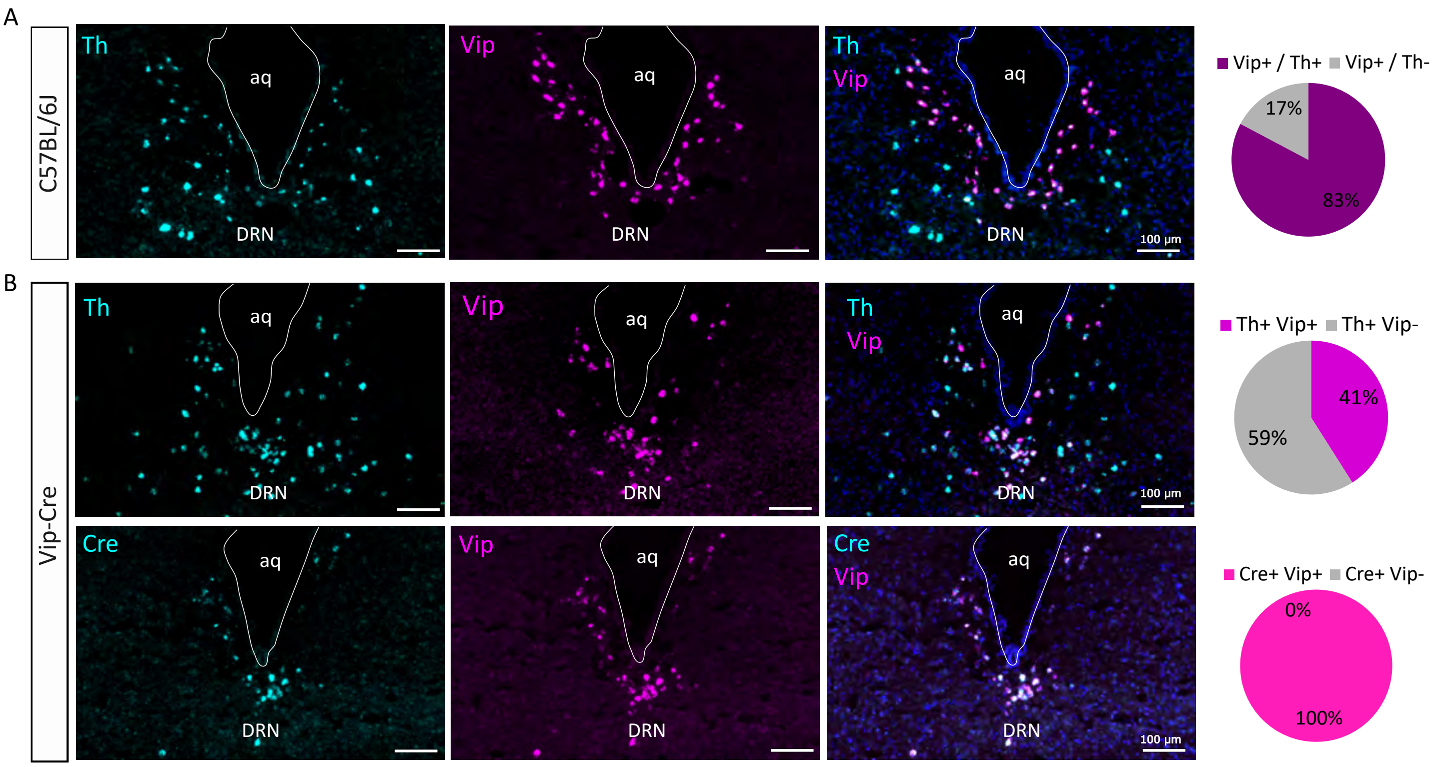

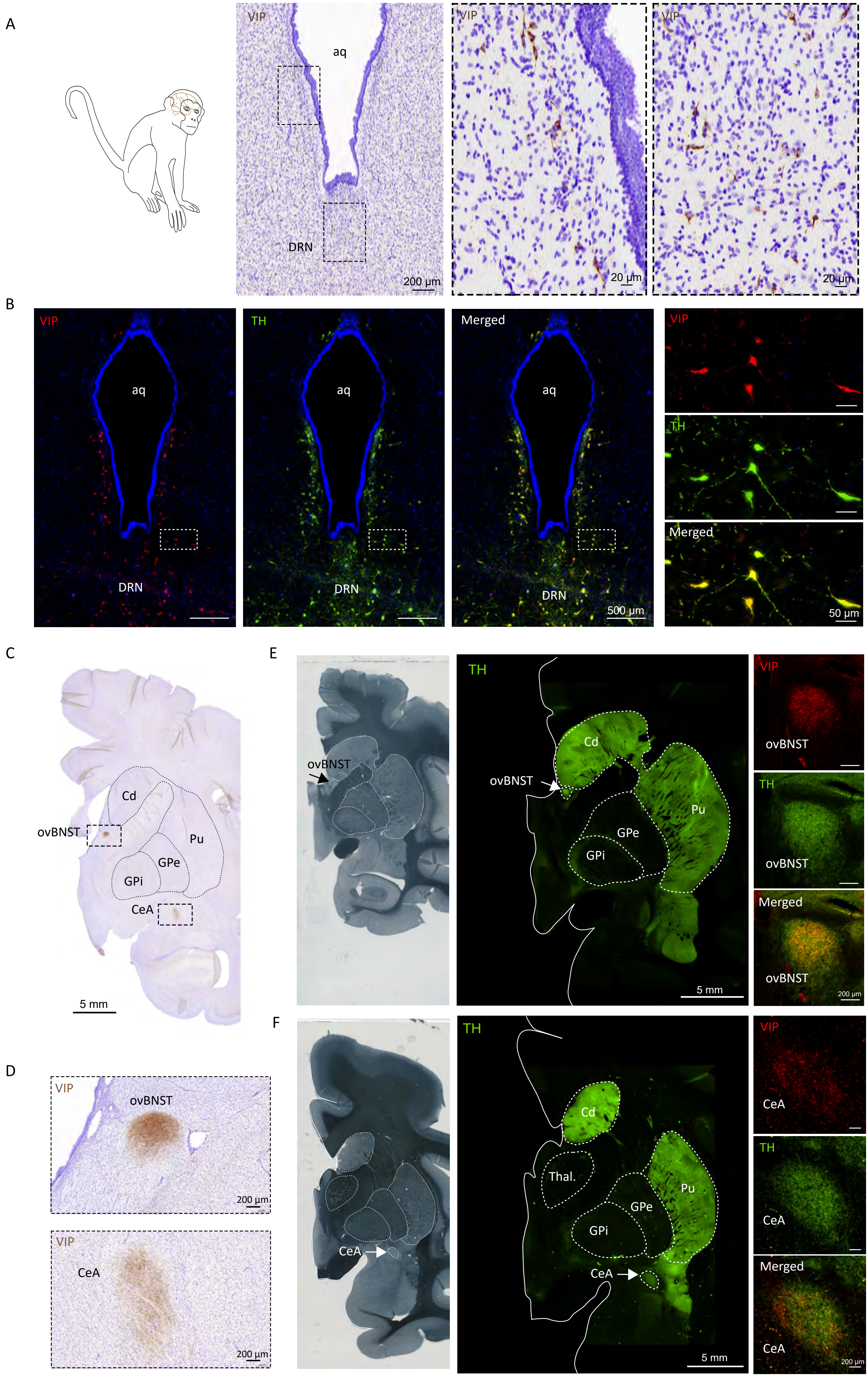

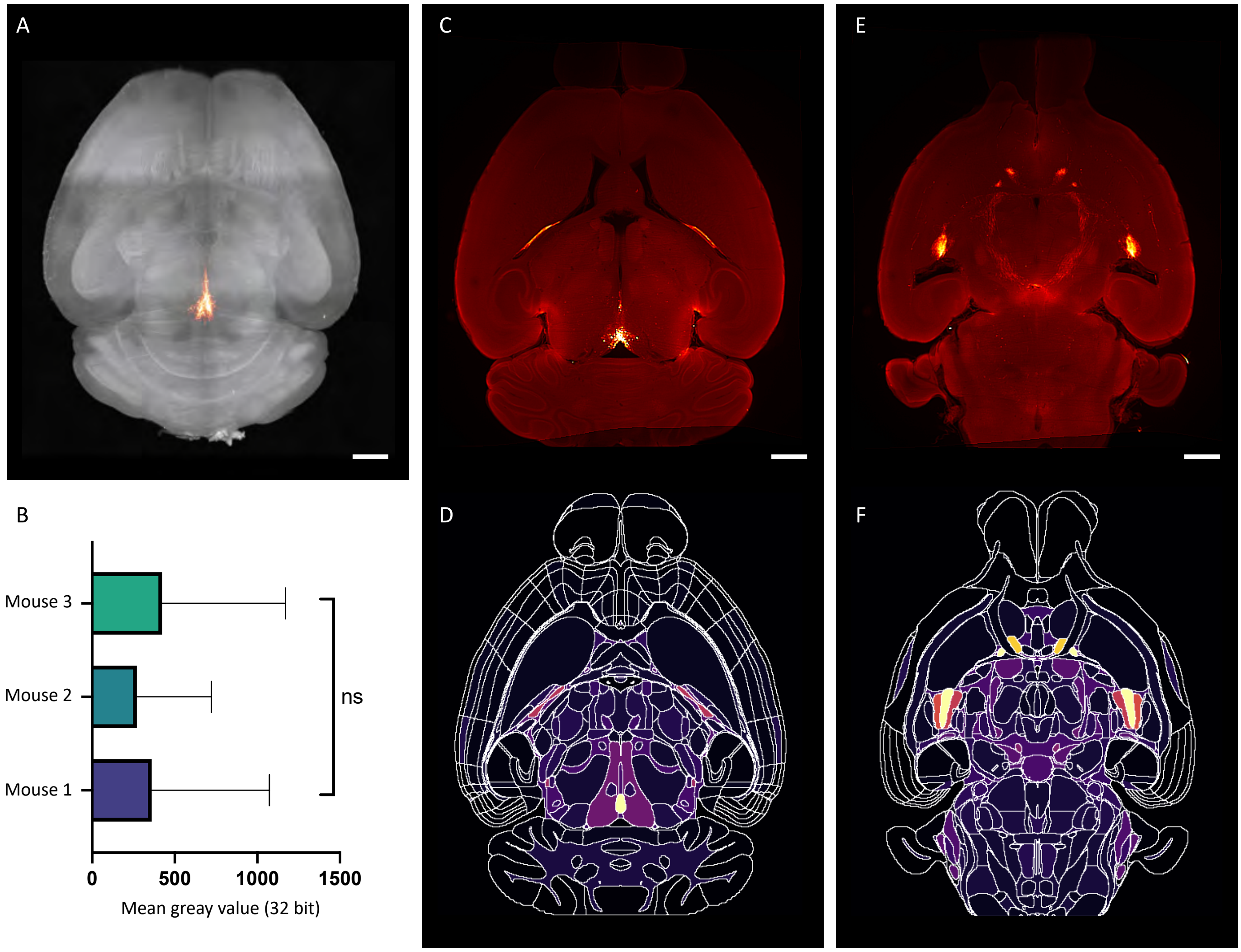

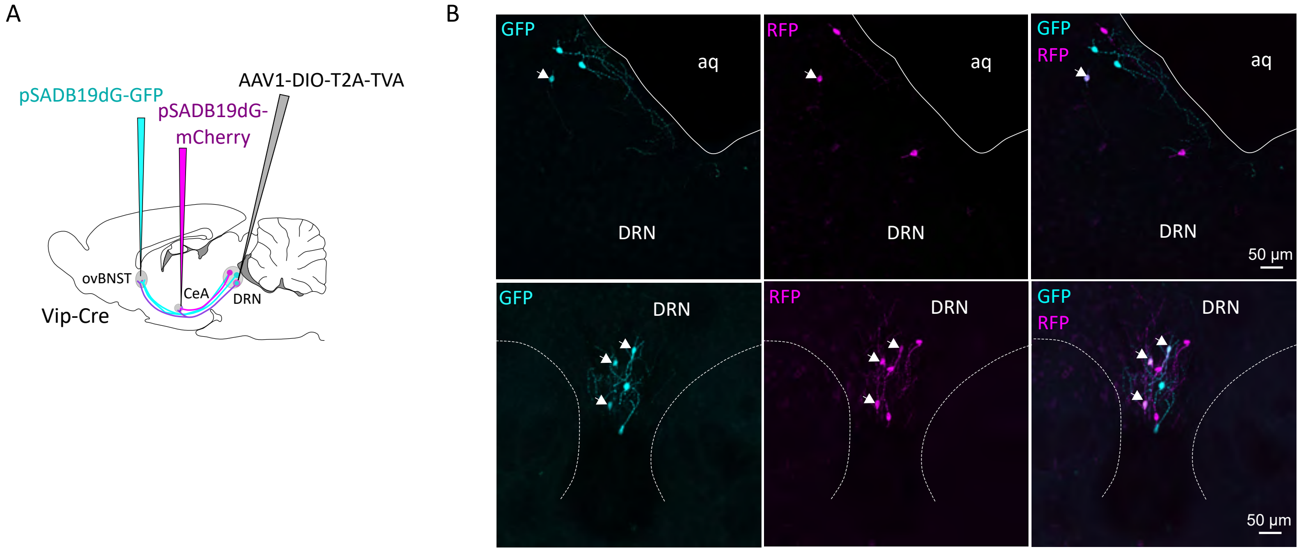

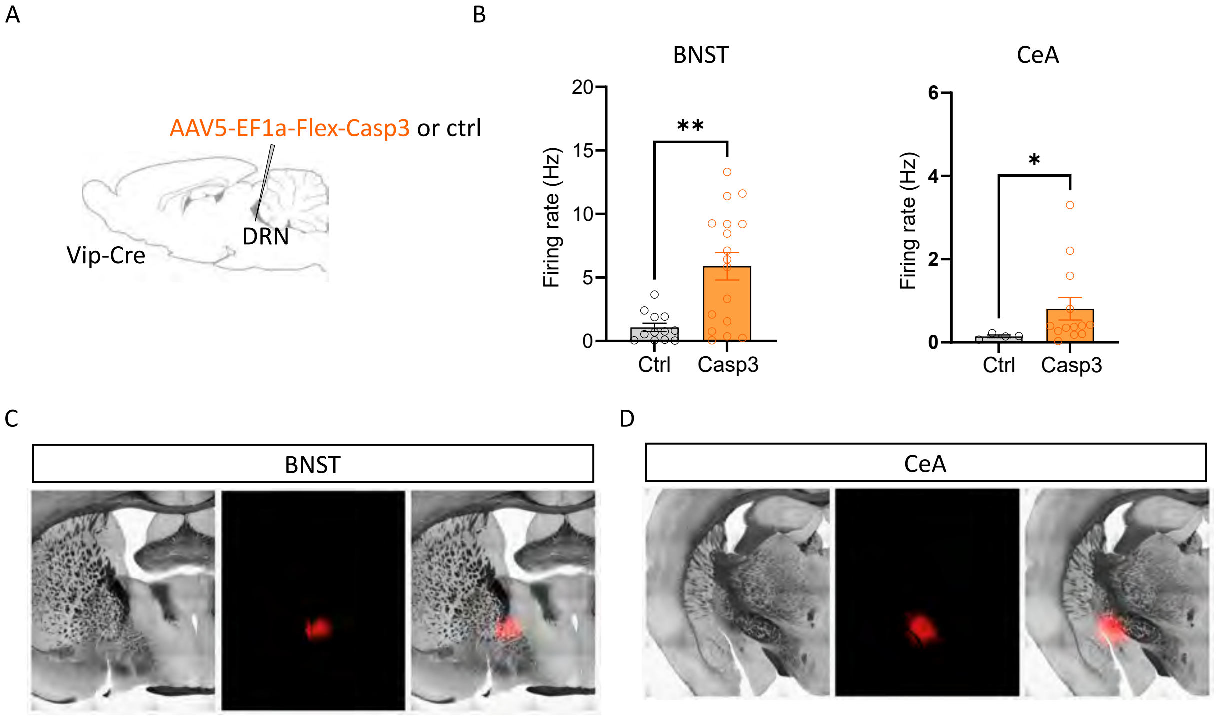

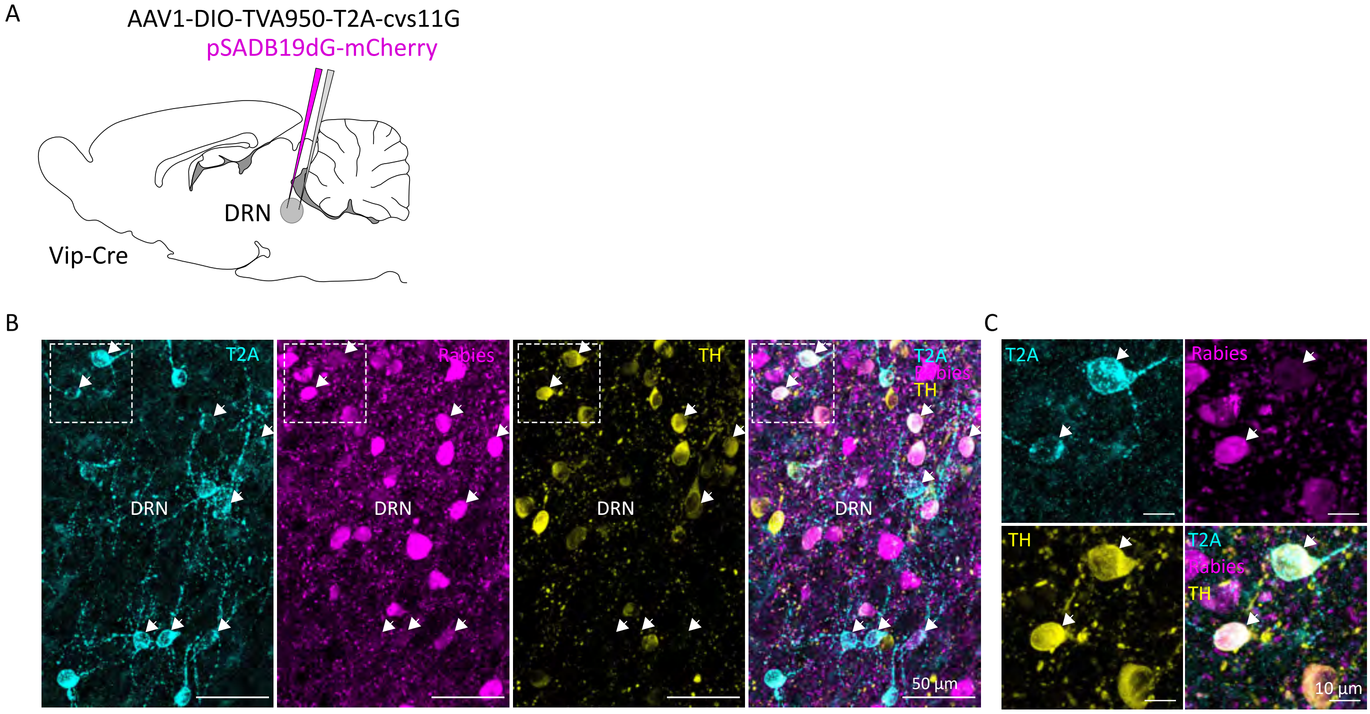

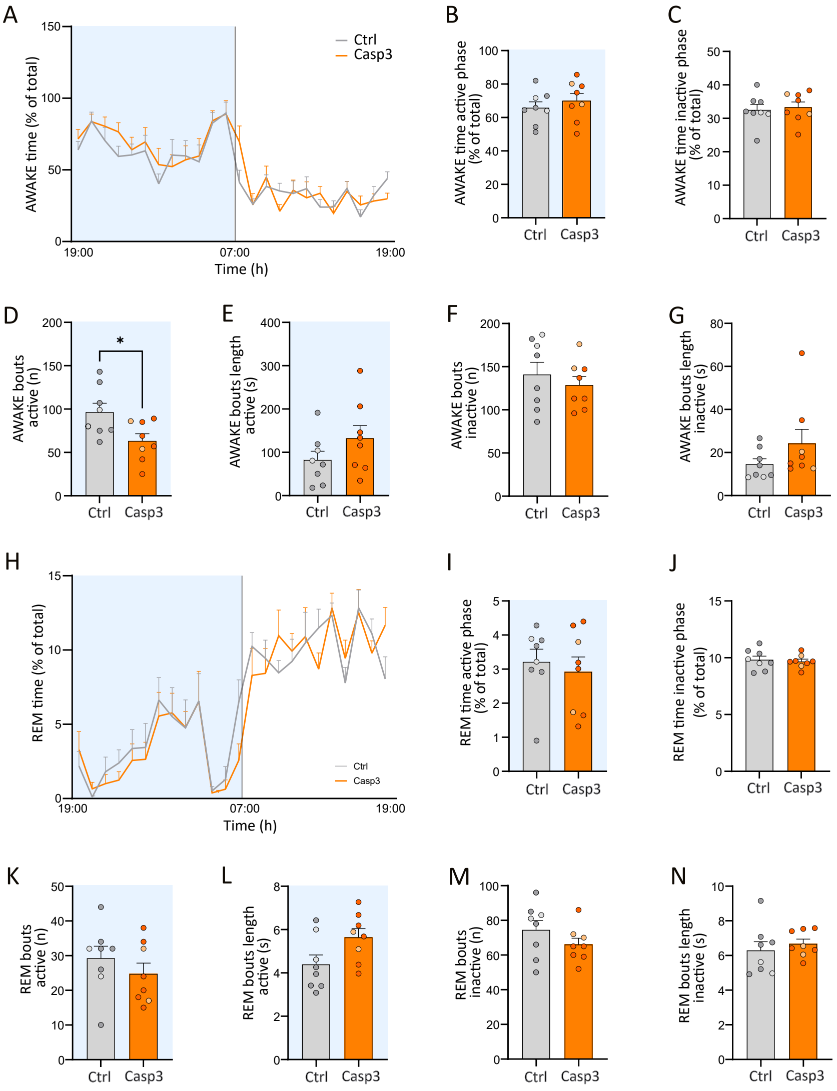
